# Cep57 and Cep57L1 cooperatively maintain centriole engagement during interphase to ensure proper centriole duplication cycle

**DOI:** 10.1101/2020.05.10.086702

**Authors:** Kei K. Ito, Koki Watanabe, Haruki Ishida, Kyohei Matsuhashi, Takumi Chinen, Shoji Hata, Daiju Kitagawa

**Affiliations:** Department of Physiological Chemistry, Graduate school of Pharmaceutical Science, The University of Tokyo, Bunkyo, Tokyo 113-0033, Japan

**Keywords:** Centriole engagement, Centriole disengagement, Centriole, Centrosome, Chromosome segregation, Cep57, Cep57L1, Plk1

## Abstract

Centrioles duplicate in the interphase only once per cell cycle. Newly formed centrioles remain associated with their mother centrioles. The two centrioles disengage at the end of mitosis, which licenses centriole duplication in the next cell cycle. Therefore, timely centriole disengagement is critical for the proper centriole duplication cycle. However, the mechanisms underlying centriole engagement during interphase are poorly understood. Here, we show that Cep57 and Cep57L1 cooperatively maintain centriole engagement during interphase. Co-depletion of Cep57 and Cep57L1 induces precocious centriole disengagement in the interphase without compromising cell cycle progression. The disengaged daughter centrioles convert into centrosomes during interphase in a Plk1-dependent manner. Furthermore, the centrioles reduplicate and the centriole number increases, which results in chromosome segregation errors. Overall, these findings demonstrate that the maintenance of centriole engagement by Cep57 and Cep57L1 during interphase is crucial for the tight control of centriole copy number and thus for proper chromosome segregation.

## INTRODUCTION

The centrosome is an organelle that serves as a major microtubule organizing center (MTOC) in animal cells(Conduit et al., 2015). In mitosis, the two centrosomes migrate to opposite sides of the cell and facilitate the formation of a bipolar spindle(Petry, 2016). Therefore, the number of centrosomes must be strictly controlled for proper chromosome segregation(Nigg and Holland, 2018). Abnormalities in centrosome number cause improper spindle formation, chromosome instability, and various disorders including cancer and congenital abnormalities such as microcephaly(Bettencourt-Dias et al., 2011). After cell division, each daughter cell harbors two centrioles, and a new daughter centriole forms in proximity to the mother centriole during the S phase(Nigg and Holland, 2018). As the number of centrioles are halved after cell division, the centrioles are duplicated only once per cell cycle, ensuring that the correct number of centrioles is maintained(Gönczy and Hatzopoulos, 2019). Defects in the centriole duplication cycle can lead to aberrations in centrosome number(Nigg and Holland, 2018).

The centrosome is composed of one or two centrioles and the surrounding pericentriolar material (PCM)(Conduit et al., 2015). The centrioles are duplicated only once in the S phase(Fu et al., 2015). Each newly formed daughter centriole grows at the proximity of the mother centriole during centriole duplication and is orthogonally engaged with the mother centriole until late mitosis (centriole engagement). After mitotic exit, mother and daughter centrioles are dissociated (centriole disengagement), and both centrioles are licensed to duplicate in the next cell cycle(Tsou et al., 2009). When centriole disengagement occurs precociously in the interphase, centrioles are reduplicated within the same cell cycle(Lončarek et al., 2010; Martino et al., 2015). Such centriole reduplication results in an increase in the number of centrioles in cycling cells and may lead to chromosomal instability and a failure of cell division(Holmes et al., 2010). Thus, the maintenance of centriole engagement is one of the mechanisms limiting centriole duplication to once per cell cycle and controlling proper centrosome cycle progression. However, the molecular mechanisms underlying centriole engagement and disengagement remain largely unknown.

Recently, it has been suggested that the expanded PCM surrounds the pair of centrioles and maintains centriole engagement during mitosis(Seo et al., 2015). In particular, in human cells, pericentrin (PCNT), a major PCM scaffold protein, has been shown to be a critical factor for centriole engagement during mitosis(Lee and Rhee, 2012; Matsuo et al., 2012). PCNT is an elongated molecule that is radially oriented, with its C-terminus region near the centriole and its N-terminus extending outward to the periphery(Lawo et al., 2012; Mennella et al., 2012). A radial array of PCNT acting as a scaffold for PCM facilitates the recruitment of other PCM proteins during PCM expansion, and the depletion of PCNT causes precocious centriole disengagement in early mitosis(Matsuo et al., 2012). Importantly, we recently identified Cep57 (centrosomal protein of 57 kDa) as a binding partner of PCNT(Watanabe et al., 2019). Cep57 localizes at the vicinity of centrioles and binds to the PACT domain, a conserved C-terminus domain, of PCNT. Depletion of Cep57 perturbs the Cep57-PCNT interaction and thereby affects PCM organization in early mitosis, leading to precocious centriole disengagement(Watanabe et al., 2019). For the centriole disengagement that normally occurs at the end of mitosis, separase-dependent cleavage of PCNT, which presumably occurs around the metaphase-to-anaphase transition, is required for the disassembly of expanded PCM and subsequent centriole disengagement(Lee and Rhee, 2012; Matsuo et al., 2012). Plk1-dependent phosphorylation of PCNT has also been reported as a priming step for the separase-dependent cleavage of PCNT(Kim et al., 2015). In the prolonged G2/M phase induced by treatment with a Cdk1 inhibitor or DNA-damaging reagents, Plk1 and separase are aberrantly activated, which subsequently induces precocious centriole disengagement(Douthwright and Sluder, 2014; Inanç et al., 2010; Prosser et al., 2012). However, in contrast to the recent progress in understanding mechanisms of centriole disengagement in mitosis, very little is known about mechanisms maintaining centriole engagement during interphase.

*Centrosomal protein 57 kDa-like protein 1* (*Cep57L1*) is a paralog of the *Cep57* gene and is conserved in vertebrates. Cep57L1 was originally named for its homology to Cep57. It has been reported that the reduction of Cep57L1 expression level is responsible for the congenital absence of the anterior cruciate ligament (ACL) and the posterior cruciate ligament (PCL) in the knee(Liu et al., 2015). Although we have recently demonstrated that Cep57 localizes at the centrosome and is required for the maintenance of centriole engagement in early mitosis(Watanabe et al., 2019),neither the exact function nor localization of Cep57L1 has been reported so far.

In this study, we reveal that Cep57 and Cep57L1 redundantly regulate centriole engagement in the interphase. Co-depletion of Cep57 and Cep57L1 causes precocious centriole disengagement during interphase in human cells. Such precociously disengaged daughter centrioles acquire PCM and MTOC activity. The precocious centriole disengagement in interphase is accompanied by centriole reduplication, and the number of centrioles per cell gradually increases with each passing cell division, since the amplified centrioles are inevitably inherited by the daughter cells. Furthermore, the amplified centrioles cause a higher frequency of chromosome segregation errors. It is therefore most likely that defects in centriole engagement in interphase are more deleterious than those in early mitosis. Overall, these findings shed light on the molecules involved in the maintenance of centriole engagement during interphase and clarify the effects of the disruption of centriole engagement on the fidelity of chromosome segregation and cell division.

## RESULTS

### Co-depletion of Cep57 and Cep57L1 causes an increase in the number of centrosomes during interphase

We recently reported that the Cep57-PCNT interaction is crucial for the maintenance of centriole engagement during mitosis and that depletion of Cep57 causes precocious centriole disengagement in mitosis, but not in interphase (Mitosis: 42.2 ± 5.1%, Interphase: 0%, N=30 and 50, respectively, from three independent experiments) (Figures 1A and 1B)(Watanabe et al., 2019). While the mechanism of centriole engagement in mitosis is gradually being elucidated, that of interphase remains completely unknown. To address this, we sought to identify the molecules required for centriole engagement during interphase. Given that comprehensive RNAi-based screens have failed to identify such molecules thus far, we assumed that two or more molecules have redundant functions in the maintenance of centriole engagement during interphase. Taking this possibility into account, we considered Cep57 and PCNT as potential targets. We also focused on an uncharacterized paralog of Cep57 in addition to Cep57 and PCNT: Cep57L1. Cep57L1 is a conserved protein in vertebrates and consists of 460 amino acid residues with 39% sequence homology to Cep57 in *Homo sapiens* (Figure S1A). We found that depletion of Cep57L1 did not affect centriole engagement in either interphase or mitosis (Figures 1E and 1F). We accordingly tested co-depletion of two of the three proteins, Cep57, Cep57L1, and PCNT, by treating human cells with two distinct siRNAs, and observed the centrosome and centriole behaviors. Among these combinations, we first found that co-depletion of Cep57 and Cep57L1 (Cep57/Cep57L1) caused an increase in the number of centrosomes, marked by Cep192, in interphase (46.6 ± 17.7%, N=50, from three independent experiments) (Figures 1C and 1D). Interestingly, we next found by using a centriole maker, CP110, that approximately 20% of the Cep57/Cep57L1-depleted cells exhibited precocious centriole disengagement in the interphase (16.0 ± 5.3%, N=50, from three independent experiments, we defined precocious centriole disengagement when at least one centriole is more than 0.75 μm away from the others) (Figures 1C and 1E) and that approximately 10% of the cells had more than four centrioles (11.3 ± 7.6%) (Figures 1C and 1F). These phenotypes provoked by the depletion of Cep57/Cep57L1 were also confirmed by using a different siRNA and other human cell lines (Figures S1B and S1C). As expected, this phenotype was rescued by expressing a synthetic RNAi-resistant Cep57 or Cep57L1 construct (siControl: 2.2 ± 1.9%, siCep57/Cep57L1: 33.2 ± 2.9%, siCep57/Cep57L1 + Cep57 expression: 10.0 ± 3.3%, siCep57/Cep57L1 + Cep57L1 expression: 6.7 ± 3.3%, N=30, from three independent experiments) (Figures 1G and 1H). Taking these results together, we conclude that the co-depletion of Cep57 and Cep57L1 causes the aberrations in the number of centrosomes and centrioles during interphase

**Figure 1.**
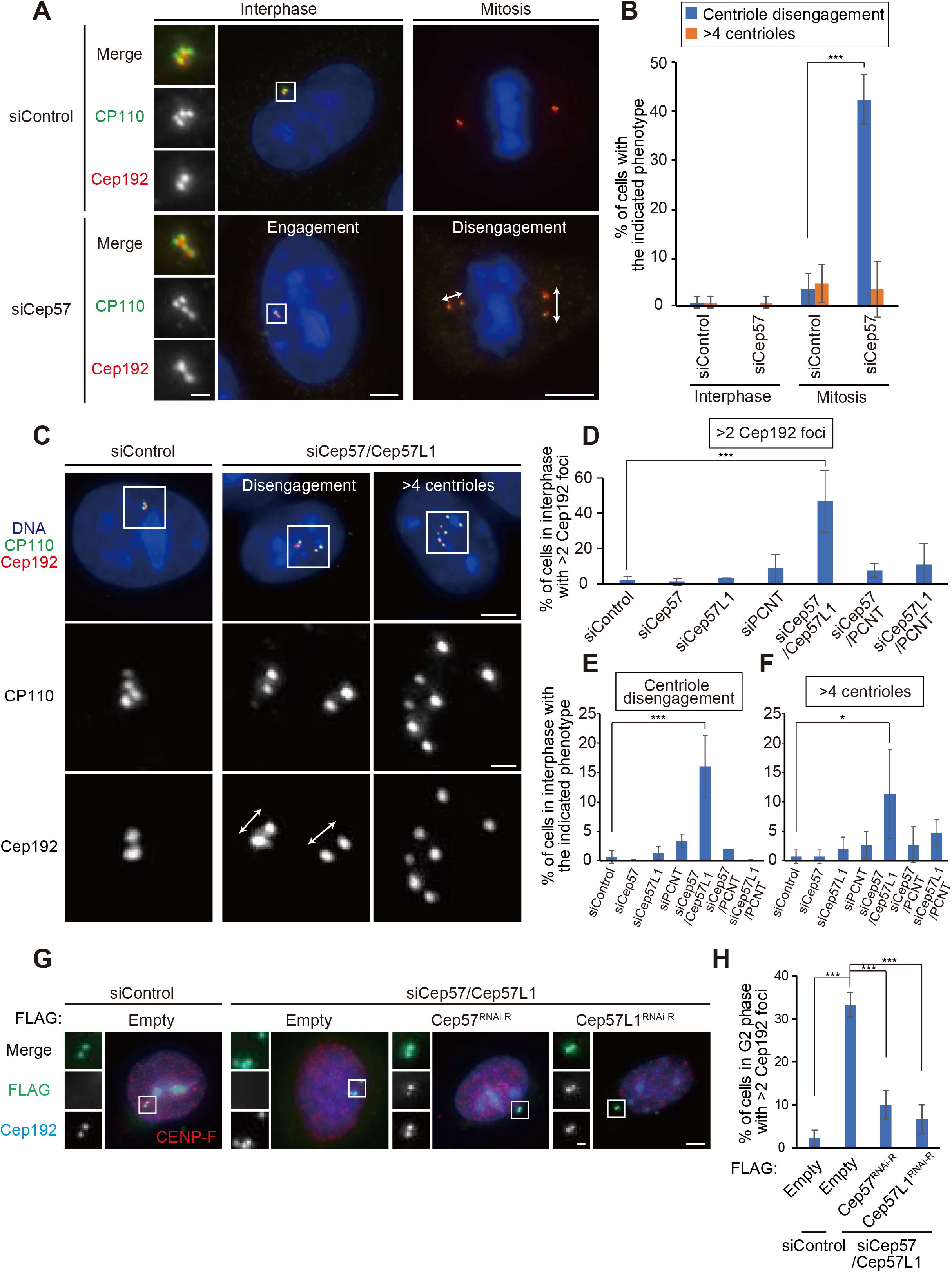
Co-depletion of Cep57 and Cep57L1 causes the increase of centrosome number during interphase. (**A**) Depletion of Cep57 induced precocious centriole disengagement in mitosis, but not in interphase. HeLa cells were treated with siControl or siCep57, and immunostained with antibodies against CP110 (green) and Cep192 (red). (**B**) Histograms represent frequency of cells in interphase and mitosis with the indicated phenotypes observed in (A). Values are mean percentages ± s.d. from three independent experiments (n = 50 for interphase, 30 for mitosis for each experiment). (**C**) Co-depletion of Cep57 and Cep57L1 (Cep57/Cep57L1) induced precocious centriole disengagement and increase of centrosome number in interphase. HeLa cells were treated with siControl or siCep57/Cep57L1, and immunostained with antibodies against CP110 (green) and Cep192 (red). (**D-F**) Histograms represent frequency of the interphase cells with >2 Cep192 foci (in (D)), centriole disengagement (in (E)) or >4 centrioles (in (F)), respectively. Values are mean percentages ± s.d. from three independent experiments. (n = 50 for each experiment). (**G**) Precocious centriole disengagement in interphase induced by Cep57/Cep57L1 co-depletion was rescued by exogenous expression of Cep57 or Cep57L1. HeLa cells were treated with siControl or siCep57/Cep57L1, followed by the transfection with FLAG empty (control), FLAG-Cep57 (RNAi-resistant) or FLAG-Cep57L1 (RNAi-resistant). The cells were immunostained with antibodies against FLAG (green), CENP-F (red) and Cep192 (cyan). (**H**) Histograms represent frequency of cells in the G2 phase with >2 Cep192 foci in each condition observed in (G). Values are percentages from three independent experiments (n = 30 for each experiment). All scale bars, 5 μm in the low-magnified view, 1 μm in the inset. Two-tailed, unpaired Welch’s t-test was used in (A) to obtain p value. Dunnett’s multiple comparisons test was used in (D), (E), (F) and (H) to obtain p value. ***p < 0.001; *p < 0.05.

### Cep57 and Cep57L1 redundantly regulate centriole engagement during interphase

In general, the presence of four separate centrioles in the interphase can stem from a failure of cytokinesis. To investigate whether the phenotype seen with co-depletion of Cep57 and Cep57L1 is due to a failure of cytokinesis or precocious centriole disengagement during interphase, we immunostained HeLa cells with an antibody against ODF2, a marker of old mother centrioles (Figures S2A and S2B). If the four separate centrioles were the consequence of a failure of cytokinesis, the number of old mother centrioles in an interphase cell should be two. However, more than 80% of the HeLa cells with four separate centrioles or amplified centrioles possessed only one ODF2 focus (84.8 ± 1.7%, N > 50, from two independent experiments), as was the case in control cells (100%, Figures S2A and S2B). We therefore reasoned that the separate centrioles likely resulted from precocious centriole disengagement in interphase, rather than cytokinesis failure. To confirm this idea, we determined when in the cell cycle the phenotype could be observed in Cep57/Cep57L1-depleted cells using 5-ethynyl-2’-deoxyuridine (EdU, an S phase marker) and an antibody against CENP-F (a G2 phase marker). Immunofluorescence with the cell cycle markers indicated that, as in control cells, there were two centrioles in the G1 phase in Cep57/Cep57L1-depleted cells, (EdU-negative, CENP-F-negative cells), but approximately 20% of S phase cells (EdU-positive, CENP-F-negative cells) and 40% of G2 phase cells (CENP-F-positive cells) exhibited four or more separate centrioles (S phase: 24.9 ± 2.0%, G2 phase: 42.9 ± 10.1%, N=30, from three independent experiments) (Figures 2A, 2B and 2C). These data strongly suggest that the presence of the four separate centrioles observed during interphase in Cep57/Cep57L1-depleted cells stem from precocious centriole disengagement in the S and G2 phases. Furthermore, live-imaging analysis using HeLa cells stably expressing GFP-centrin1 (HeLa-GFP-centrin1) confirmed that precocious disengagement occurred during interphase in Cep57/Cep57L1-depleted cells (Figure 2D and Supplementary Movie 1). We also measured cumulative percentages of the disengagement phenotype based on live-imaging data and found that approximately 60% of Cep57/Cep57L1-depleted cells (16 out of 26 cells) exhibited a disengagement phenotype before cell round-up, that is, in the interphase (Figure 2E, Mean time: siControl t=4.20 h, siCep57/Cep57L1 t=-2.76 h, Time zero corresponds to the cell round-up). Importantly, we also found that centriole disengagement was occasionally followed by centriole reduplication (Figure 2E), and that around one-fifth of Cep57/Cep57L1-depleted cells possessed more than four centrioles in the G2 phase (22.1 ± 11.1%, N=30, from three independent experiments) (Figure 2C). Taken together, these data suggest that Cep57 and Cep57L1 cooperatively regulate the maintenance of centriole engagement during interphase and thus suppress centriole reduplication within the same cell cycle.

**Figure 2.**
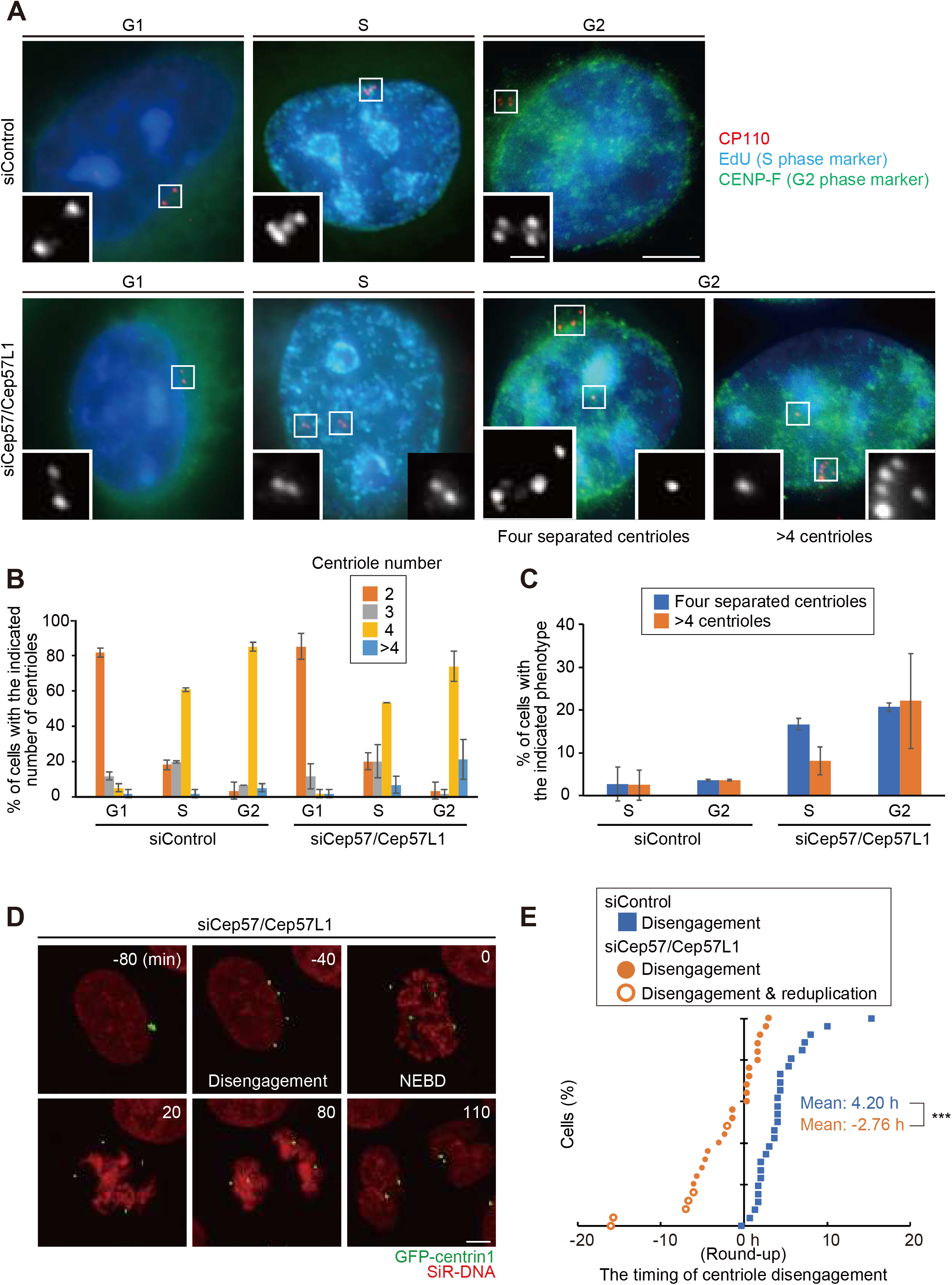
Cep57 and Cep57L1 redundantly regulate centriole engagement during interphase. (**A**) Cep57/Cep57L1 co-depletion induced four separated centrioles in the G2 phase. HeLa cells were treated with siControl or siCep57/Cep57L1 for 36 h in the presence of 5-ethynyl-2’-deoxyuridine (EdU, S phase marker, cyan) for the last 30 min before fixation, and immunostained with antibodies against CP110 (red) and CENP-F (green). Scale bars, 5 μm in the low-magnified view, 1 μm in the inset. (**B**) Histograms represent the number of centrioles in the G1, S and G2 phase treated with the indicated siRNAs in (A). (**C**) Histograms represent frequency of cells in the S and G2 phase treated with the indicated siRNAs exhibiting the indicated phenotype. Values are mean percentages ± s.d. from three independent experiments (n = 30 for each experiment) in (B) and (C). (**D**) Time-lapse observation of cells upon Cep57/Cep57L1 depletion. HeLa cells stably expressing GFP-centrin1 (HeLa-GFP-centrin1) were treated with siControl or siCep57/Cep57L1 in the presence of SiR-DNA (200 nM). Scale bar, 5 μm. (**E**) Cumulative scatter plot indicates the duration from cell round-up to centriole disengagement observed in (D). Orange open circles indicate precocious centriole disengagement accompanied by centriole reduplication. Two-tailed, unpaired Welch’s t-test was used in (E) to obtain p value. ***p < 0.001.

Previous studies have reported that the G2 phase cell-cycle arrest induced by treatment with a Cdk1 inhibitor or DNA-damaging reagents caused precocious centriole disengagement(Douthwright and Sluder, 2014; Prosser et al., 2012). Therefore, to examine whether co-depletion of Cep57 and Cep57L1 induces cell cycle arrest in the interphase, we performed a live-imaging analysis and measured the duration of the period from anaphase onset to round-up in the next mitosis. In Cep57/Cep57L1-depleted cells, the duration was not significantly altered compared with control cells (siControl: 26.8 ± 2.2 h, siCep57/Cep57L1: 26.2 ± 3.6 h, N=30) (Figure S2B). In addition, FACS profiling analysis revealed that the distribution of cell cycle phases was not affected by Cep57/Cep57L1 depletion (Figure S2C). Overall, these findings reveal that the co-depletion of Cep57 and Cep57L1 causes precocious centriole disengagement starting in the S phase without affecting the cell cycle progression

### Precociously disengaged daughter centrioles are converted to centrosomes in the G2 phase

Given that after depletion of Cep57/Cep57L1 precocious centriole disengagement was apparent starting in the S phase, and that centriole formation proceeds during the S phase, we next examined whether the disengaged daughter centrioles were fully elongated. To address this, we used POC5, which is known to be incorporated at the final stage of new centriole formation(Azimzadeh et al., 2009; Chang et al., 2016). Immunostaining with an antibody against POC5 revealed that approximately half of the disengaged centrioles incorporated POC5 (57.1% ± 4.0%, N=30, from three independent experiments), suggesting that upon depletion of Cep57/Cep57L1, a daughter centriole can disengage from its mother centriole during the process of its formation (Figures S3A and S3B).

Since fully matured daughter centrioles are converted into centrosomes after mitosis (a process called centriole-to-centrosome conversion)(Wang et al., 2011), we then asked if the precociously disengaged daughter centrioles accomplish the centriole-to-centrosome conversion in the interphase. In normal cells, around mitotic exit, a daughter centriole becomes a functional centrosome and recruits PCM components and centriole duplication factors, such as PCNT and Cep152, respectively. For this conversion, procentriole formation requires the centriolar recruitment of Cep295 and Cep192(Fu et al., 2016; Tsuchiya et al., 2016). Similar to Cep192, Cep295 was localized at almost all of the precociously disengaged daughter centrioles (Figure S3C). In contrast, the disengaged daughter centrioles did not always acquire PCM components (PCNT), suggesting that PCM proteins were gradually recruited to the disengaged daughter centrioles. Indeed, Cep57/Cep57L1-depleted cells with more than two PCNT-positive centrioles were relatively rare before the G2 phase (15.0 ± 2.4%, N=30 from three independent experiments), but gradually increased during the G2 phase (51.7 ± 2.4%) and were most frequently observed after the G2 phase was completed (mitosis, 85.0 ± 11.8%) (Figures S4A and S4B). This observation shows that the precociously disengaged daughter centrioles gradually recruited PCM components to their surroundings mainly during the G2 phase and mitosis (Figure S4C). Furthermore, such disengaged daughter centrioles nucleated microtubules after depolymerization of microtubules by cold treatment, indicating that these centrioles had acquired MTOC activity during interphase (Figure S4D). Taken together, these results indicate that upon Cep57/Cep57L1 depletion, the precociously disengaged centrioles can be converted to centrosomes starting in the interphase

### Co-depletion of Cep57 and Cep57L1 induces centriole disengagement in the interphase even without Plk1 kinase activity

To gain insight into the mechanisms by which Cep57/Cep57L1 depletion causes precocious centriole disengagement in the interphase, we tested a requirement of the known factors involved in centriole disengagement at the end of mitosis. Previous studies have reported that canonical centriole disengagement requires Plk1 activity, and that the inhibition of Plk1 perturbs centriole disengagement at the end of mitosis (Tsou et al., 2009). Precocious centriole disengagement in G2/M phase-arrested cells was also suppressed by Plk1 inhibition (Prosser et al., 2012). To determine whether the precocious centriole disengagement in Cep57/Cep57L1-depleted cells also requires Plk1 activity, we treated G2 phase-arrested cells or Cep57/Cep57L1-depleted cells with a small molecular inhibitor of Plk1 (BI2536). As expected, the precocious centriole disengagement in G2/M-arrested cells was suppressed by BI2536 treatment (Figures 3A and 3B). In contrast, BI2536 treatment did not suppress the precocious centriole disengagement or the increase in the number of centrioles in Cep57/Cep57L1-depleted cells (DMSO: 46.7 ± 10.0%, BI2536: 45.6 ± 10.1%, N=30 from three independent experiments) (Figures 3C and 3D). These data indicate that the precocious centriole disengagement that occurs in Cep57/Cep57L1-depleted cells does not depend on Plk1 activity, unlike canonical centriole disengagement or cell-cycle-arrest-induced precocious centriole disengagement

**Figure 3.**
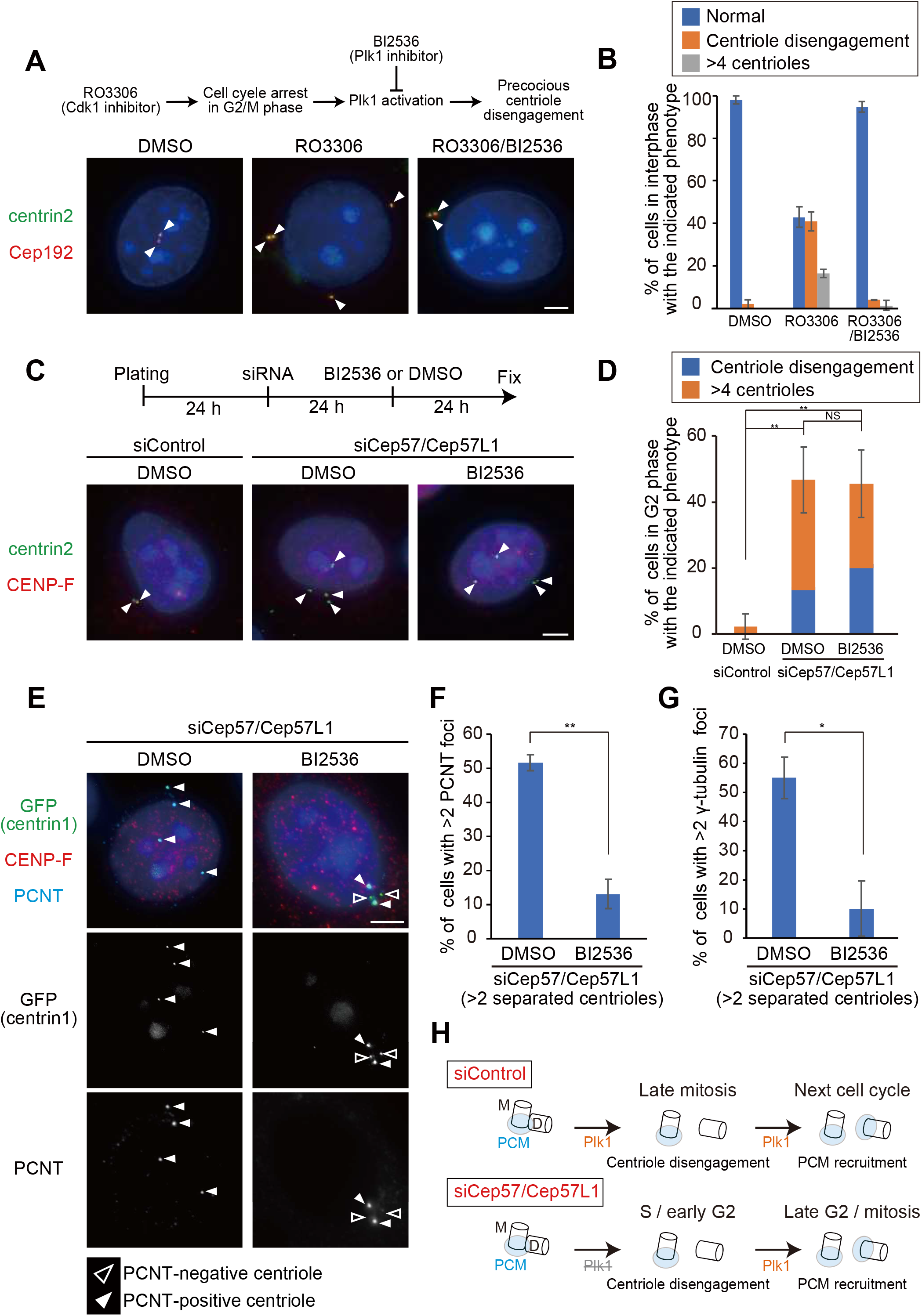
Inhibition of Plk1 prevents disengaged daughter centrioles from acquiring PCM, but does not suppress precocious centriole disengagement. (**A**) Precocious centriole disengagement caused by G2/M phase arrest was suppressed by BI2536 (Plk1 inhibitor, 100 nM). HeLa cells were synchronized in the G1/S phase by Aphidicolin (1.2 μg/mL) for 17 h, then released into fresh medium for 4 h. Next, the cells were treated with DMSO (control), RO3306 (Cdk1 inhibitor, 10 μM), or both of RO3306 and BI2536 (100 nM) for 24 h. HeLa cells were immunostained with antibodies against centrin2 (green) and Cep192 (red). Scale bar, 5 μm. (**B**) Histograms represent frequency of the interphase cells with the indicated phenotype observed in (A). Values are mean percentages ± s.d. from three experiments (n = 50 for each experiment). (**C**) Precocious centriole disengagement in Cep57/Cep57L1-depleted cells was not suppressed by BI2536. HeLa cells were treated with siControl or siCep57/Cep57L1 for 24 h, followed by treatment of DMSO (control) or BI2536 (100 nM) for additional 24 h. The cells were immunostained with antibodies against centrin2 (green) and CENP-F (red). (**D**) Histograms represent frequency of cells in the G2 phase with the indicated phenotype observed in (C). Values are mean percentages ± s.d. from three experiments (n = 30 for each experiment). (**E**) BI2536 suppressed recruitment of PCNT at disengaged daughter centrioles. HeLa-GFP-centrin1 cells were treated as in (C), and immunostained with antibodies against GFP (green), CENP-F (red) and PCNT (cyan). White/black arrowheads indicate PCNT positive/negative centrioles, respectively. (**F, G**) Histograms represent frequency of cells with >2 PCNT or γ-tubulin foci among the G2 phase cells with disengaged or amplified centrioles. Values are mean percentages ± s.d. from three experiments (n = 30 for each experiment). (**H**) Schematic illustration of the results in Figure 3. The activity of Plk1 is required for the recruitment of PCM components at the disengaged daughter centrioles, but not for precocious centriole disengagement itself. All scale bars, 5 μm. Tukey’s multiple comparisons test was used in (D) against total value of centriole disengagement and >4 centrioles to obtain p value. Two-tailed, unpaired Student’s t-test was used in (F) and (G) to obtain p value. **p < 0.01; *p < 0.05; NS not significantly different (p > 0.05).

### Plk1 activity is required for PCM recruitment to disengaged daughter centrioles, presumably in the G2 phase

Since Plk1 is necessary not only for centriole disengagement but also for daughter centrioles to acquire PCM in normal cells (Wang et al., 2011), we next asked if Plk1 activity is also required for PCM recruitment at precociously disengaged daughter centrioles in Cep57/Cep57L1-depleted cells. To address this, we treated Cep57/Cep57L1-depleted cells with BI2536 and counted the number of PCM-positive centrioles. Interestingly, in Cep57/Cep57L1-depleted cells, BI2536 treatment significantly reduced the number of cells with disengaged centrioles that have acquired PCNT (13.1 ± 4.3%, N=30, from three independent experiments), compared to the controls (51.6 ± 2.4%) (Figures 3E and 3F), indicating that Plk1 activity is required for PCNT recruitment to the disengaged daughter centrioles. Similarly, γ-tubulin recruitment to the disengaged daughter centrioles was also suppressed by BI2536 treatment (DMSO: 55.1 ± 7.1%, BI2536: 10.0 ± 9.4%, N=30, from three independent experiments) (Figure 3G). Overall, these findings demonstrate that the precociously disengaged daughter centrioles acquire PCM during the interphase in a Plk1-dependent manner (Figure 3H). Given that Plk1 is activated starting from the late G2 stage preceding mitosis (Schmucker and Sumara, 2014) and is necessary for PCM recruitment to the disengaged daughter centrioles, it is reasonable that precociously disengaged daughter centrioles gradually acquire PCNT in the G2 phase (Figures S4A and S4B)

### Cep57 and Cep57L1 have distinct properties in spite of their relatively high amino acid sequence homology

While Cep57 and Cep57L1 cooperatively regulate centriole engagement in the interphase, centriole engagement in mitosis is maintained by Cep57 but not by Cep57L1. This suggests that Cep57 and Cep57L1 share redundant and distinct functions in centriole engagement. To investigate the similarities and differences between the functions of Cep57 and Cep57L1 in the regulation of centrioles, we performed localization and domain analysis of Cep57 and Cep57L1. We first sought to compare the detailed localization pattern of Cep57 and Cep57L1 at the centrosome. To do this, we immunostained HeLa cells with an antibody against Cep57L1 and found that endogenous Cep57L1 localized at the centrosome (Figure 4A). We also noticed that the level of Cep57L1 signal at new mother centrioles gradually increased in the interphase (Figure 4A), similarly to Cep57 (Watanabe et al., 2019). Moreover, structured illumination microscopy (SIM) analysis showed that Cep57L1 formed a ring-like structure around the mother centriole (Figure 4B). The diameter of the Cep57L1 ring was 167.7 ± 35.3 nm, which is close to that of the Cep57 ring (219.9 ± 13.9 nm in (Watanabe et al., 2019)) (Figure 4B). The specificity of the antibody was validated using siRNA against Cep57L1 (Figure 4C). Interestingly, recruitment of Cep57 and Cep57L1 at the centrosome did not depend on each other. Therefore, these results indicate that Cep57 and Cep57L1 show a similar centriolar distribution, but independently localize to the centrosome.

**Figure 4.**
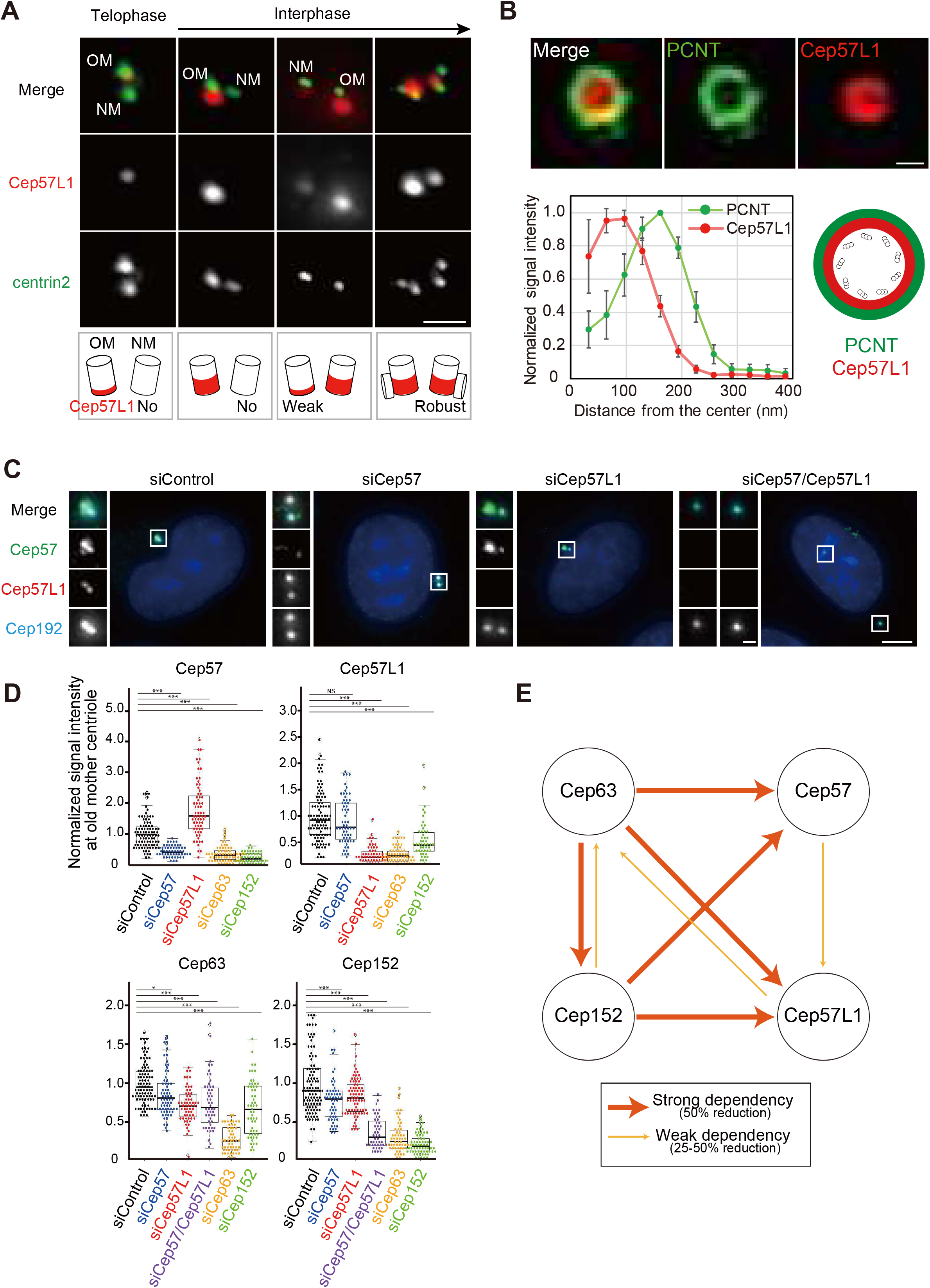
Cep57L1, a paralog of Cep57 conserved in vertebrates, shows similar centriolar distribution to Cep57. (**A**) Centriolar distribution of Cep57L1 at different cell cycle stages. HeLa cells were immunostained with antibodies against Cep57L1 (red) and centrin2 (green). Scale bar, 1 μm. (**B**) SIM images representing top view of Cep57L1 (red) and PCNT (green) at mother centrioles. Scale bar, 200 nm. The graph shows radial profiles from the center of the Cep57L1 and PCNT rings. Values are mean normalized intensity ± s.d. (n = 5). (**C**) The signals of Cep57L1 at the centrosome were attenuated by siCep57L1 or siCep57/Cep57L1, but not by siCep57. HeLa cells were treated with siControl, siCep57, siCep57L1 or both of siCep57/Cep57L1, and immunostained with antibodies against Cep57 (green), Cep57L1 (red) and Cep192 (cyan). Scale bars, 5 μm in the low-magnified view, 1 μm in the inset. (**D**) Beeswarm plots piled on boxplots represent the normalized signal intensity of Cep57, Cep57L1, Cep63 and Cep152 at the old mother centrioles upon the indicated siRNAs (n = 50). (**E**) Schematic of the dependency of the centrosome localization among Cep57, Cep57L1, Cep63 and Cep152. The wide arrows indicate strong dependency defined as more than 50% reduction of the signal intensity, and narrow arrows indicate weak dependency defined as 25-50% reduction of the signal intensity. Tukey’s multiple comparisons test was used in (D) to obtain p value. *p < 0.05; ***p < 0.001; NS not significantly different (p > 0.05).

To further examine the similarities and differences between Cep57 and Cep57L1 function, we next performed domain analysis of Cep57 and Cep57L1. We previously reported that the conserved PINC (present in N-terminus of Cep57) domain in the N-terminus of Cep57 is required for its centriolar localization and the interaction with PCNT, a functional binding partner (Watanabe et al., 2019). Since Cep57L1 also has a PINC domain, we asked if the role of the PINC motif in Cep57L1 was similar to its role in Cep57. To address this, we constructed plasmids expressing the full-length Cep57L1 or a mutant lacking the PINC motif (52-86 a.a.). Unexpectedly, immunofluorescent (IF) and co-immunoprecipitation (co-IP) analyses with the mutant revealed that the PINC motif of Cep57L1 was dispensable for the centrosome localization (Figure S5A) and for the binding to the PCNT C-terminal region, including the PACT domain (Figure S5B). These data suggest that the PINC motif has different functions between Cep57 and Cep57L1. In addition to the PINC motif, the Pfam protein family database predicted the presence of microtubule-binding domains in both Cep57 and Cep57L1 (Figure S1A). As previously reported, overexpressed Cep57 was occasionally localized on the microtubule network via its microtubule-binding domain (Figure S5C) (Momotani et al., 2008). In contrast, overexpressed Cep57L1 did not show such a localization pattern, but instead aggregated in the cytoplasm (Figure S5C). This result suggests that the putative microtubule-binding domain of Cep57L1 does not possess a binding affinity for microtubules. Furthermore, co-IP assays of Cep57 and Cep57L1 proteins in HEK293T cells detected self-interaction of Cep57L1, but not of Cep57 (Figure S5D). Overall, these findings show that Cep57 and Cep57L1 have distinct properties in spite of their relatively high amino acid sequence homology

### Interdependency of centrosome localization between Cep57, Cep57L1, Cep63, and Cep152

We next searched for the proteins recruiting Cep57L1 to the centrosome. Considering the similar distribution patterns of Cep57 and Cep57L1 at centrosomes, we postulated that the localization of Cep57 and Cep57L1 around the centriole wall is regulated by common proteins. Cep57 is recruited to the centrosome dependent on Cep63, Cep152, and NEDD1 (Aziz et al., 2018; Lukinavičius et al., 2013; Wu et al., 2012). Of these three proteins, Cep63 and Cep152 were demonstrated to form a trimeric complex with Cep57 at the centrosome (Lukinavičius et al., 2013). Consistent with previous studies (Aziz et al., 2018; Lukinavičius et al., 2013), we confirmed that Cep63 (Median: 30.6% compared to siControl, N > 50) or Cep152 depletion (19.3%) significantly decreased the signal intensity of Cep57 at the centrosome (Figures 4D and S6A). We then assumed that Cep63 and Cep152 were also responsible for centrosomal localization of Cep57L1. As expected, depletion of Cep63 (24.8%) or Cep152 (45.8%) reduced the signal intensity of Cep57L1 at the centrosomes, indicating that both Cep57 and Cep57L1 localization at the centrosomes was partially dependent on Cep63 and Cep152 (Figures 4D and S6A). On the other hand, the signal intensity of Cep63 and Cep152 at the centrosomes was only slightly affected by the depletion of Cep57 (Cep63: 81.0%, Cep152: 80.2%) or Cep57L1 (Cep63: 70.6%, Cep152: 81.0%), while co-depletion of Cep57 and Cep57L1 reduced the signal intensity of Cep152 at the centrosomes more drastically (31.1%) (Figures 4D, S6B and S6C). Moreover, as reported in previous studies (Brown et al., 2013; Kim et al., 2019; Lukinavičius et al., 2013), the signal intensity of Cep152 was attenuated by Cep63 depletion (25.8%), and vice versa (66.4%) (Figure 4D). Hence, we propose that Cep57, Cep57L1, Cep63, and Cep152 mutually influence centrosomal localization (Figure 4E)

### Precocious centriole disengagement in the interphase results in centriole reduplication and thereby accelerates numerical centrosome abnormalities

We next examined the long-term consequence of precocious centriole disengagement in Cep57/Cep57L1-depleted cells. To this end, we treated HeLa cells with siCep57/Cep57L1 for 96 h and observed the centriole number during mitosis. Immunofluorescence revealed that approximately 80% of the Cep57/Cep57L1-depleted cells had more than four centrioles during mitosis (80.1 ± 9.0%, N=50, from three independent experiments). In contrast, this was not the case for Cep57-depleted cells, in which centriole disengagement precociously occurred only in mitosis (siControl: 4.7 ± 3.6%, siCep57: 10.7% ± 1.3%) (Figures 5A and 5B). This result suggests that the precociously disengaged centrioles upon co-depletion of Cep57 and Cep57L1, were already licensed to reduplicate during the interphase within the same cell cycle. This may have thereby led to the increase in the number of centrioles. To test this idea, we monitored the number of procentrioles using an HsSAS-6 marker. The Cep57/Cep57L1-depleted cells with two pairs of engaged centrioles possessed two HsSAS-6 foci, as in control cells (siControl: 98.9 ± 1.9%, siCep57/Cep57L1: 93.3 ± 3.3%, N=30, from three independent experiments) (Figures 5C and 5D). On the other hand, more than half of the Cep57/Cep57L1-depleted cells with four separate centrioles had no HsSAS-6 foci in the interphase (60.0 ± 6.7%) (Figures 5C and 5D), suggesting that HsSAS-6 disappeared from the centrosomes, presumably just after centriole disengagement. Interestingly, in Cep57/Cep57L1-depleted cells with four HsSAS-6 foci, each HsSAS-6 focus was associated with a pair of centrioles (Figure 5C), which leads us to reason that the precociously disengaged centrioles newly acquired HsSAS-6 and reduplicated within the same cell cycle. To address this further, we performed live-cell imaging using the HeLa-GFP-centrin1 cell line, and revealed that centriole reduplication occurred after the precocious centriole disengagement in the interphase (Figure 5E). Intriguingly, we noticed that the earlier the precocious centriole disengagement occurred, the more frequently centriole reduplication could be observed (Figure 2E). In addition, live-imaging data revealed that the centrioles amplified by centriole reduplication could be inherited by the two daughter cells (Figure 5E and Supplementary Movie 2). The inherited amplified centrioles could normally duplicate in the subsequent cell cycle, which further led to numerical centrosome abnormalities. Indeed, comparing the number of centrioles between 48 h and 96 h after the siRNA treatment, the numerical abnormalities was more significant at 96 h post-treatment (Figure 5F). Overall, these findings demonstrate that precocious centriole disengagement in interphase results in numerical centrosome abnormalities with each passing cell division due to continuous centriole reduplication and the inevitable inheritance of amplified centrioles by daughter cells

**Figure 5.**
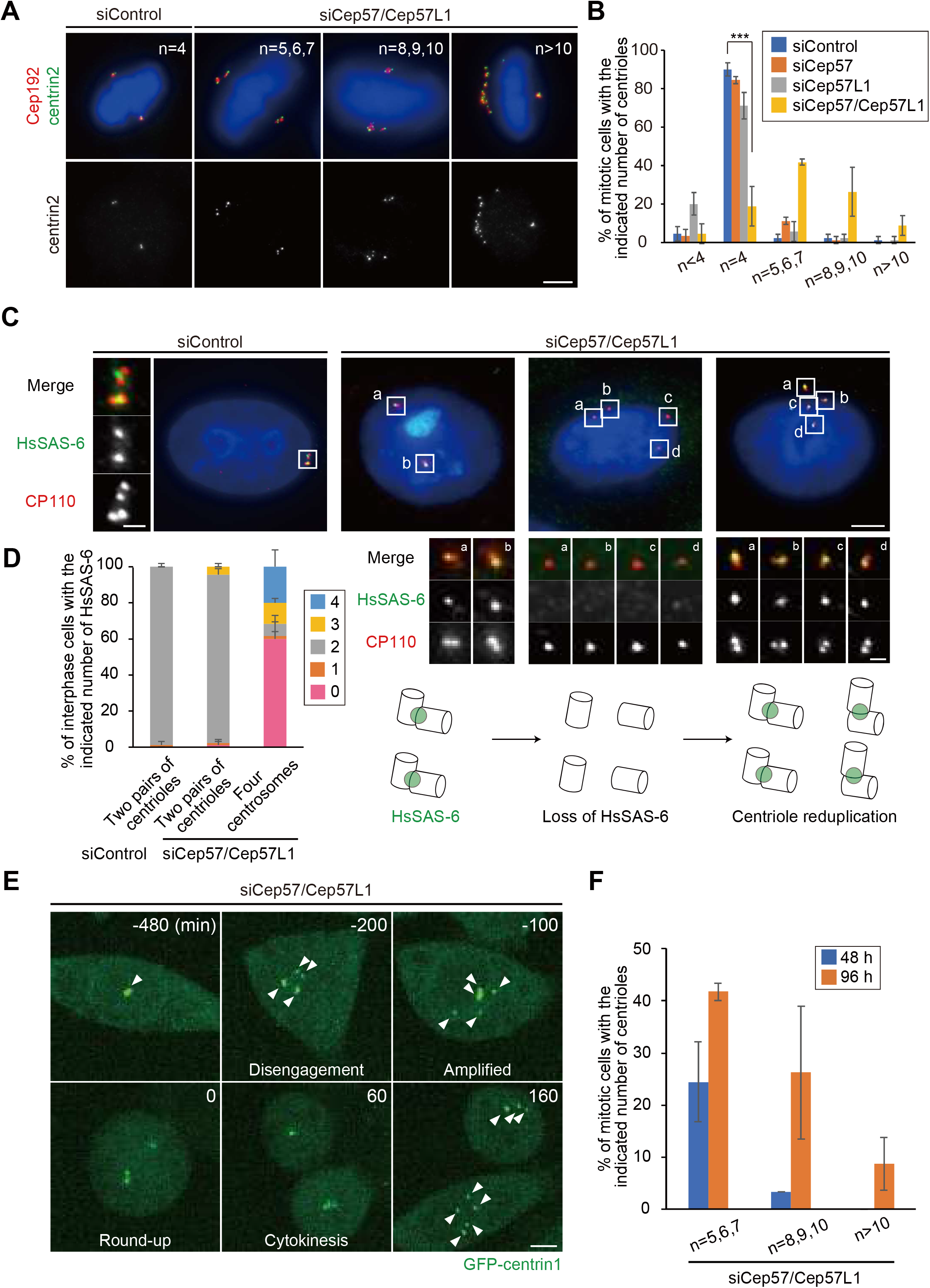
Precocious centriole disengagement results in an increase of the number of centrioles. (**A**) Long-term Cep57/Cep57L1 co-depletion increased the number of centrioles. HeLa cells were treated with siCep57/Cep57L1 for 96 h, and immunostained with antibodies against centrin2 (green) and Cep192 (red). (**B**) Histograms represent frequency of mitotic cells with the indicated number of centrioles observed in (A). Values are mean percentages ± s.d. from three independent experiments (n = 30 for each experiment). Note that depletion of Cep57L1 for 96 h slightly reduced the number of centrioles. (**C**) HsSAS-6 was absent from disengaged centrioles and was re-acquired. HeLa cells were treated with siControl or siCep57/Cep57L1, and immunostained with antibodies against HsSAS-6 (green) and CP110 (red). (**D**) Histograms represent frequency of the number of HsSAS-6 foci on the two pairs of centrioles or four centrosomes in (C). Values are mean percentages ± s.d. from three independent experiments (n = 30 for each experiment). (**E**) Precocious centriole disengagement and centriole reduplication in interphase, and inevitable inheritance of amplified centrioles to daughter cells observed in Cep57/Cep57L1-depleted cells. HeLa-GFP-centrin1 cells were treated with siCep57/Cep57L1 in the presence of SiR-DNA (200 nM). White arrowheads indicate centrosomes. (**F**) Histograms represent frequency of mitotic cells with the indicated number of centrioles 48 h or 96 h after Cep57/Cep57L1 co-depletion observed in (A) and Figure S7A. Values are mean percentages ± s.d. from three independent experiments (n = 30 for each experiment). All scale bars, 5 μm in the low-magnified view, 1 μm in the inset. Dunnett’s multiple comparisons test was used in (B) to obtain p value. ***p < 0.001.

### Centriole reduplication in Cep57/Cep57L1-depleted cells leads to the high frequency of chromosome segregation errors

Since precocious centriole disengagement causes defects in chromosome segregation (Watanabe et al., 2019), we then explored the fate of Cep57/Cep57L1-depleted cells in mitosis. We first grouped cells into four categories based on the timing of centriole disengagement and whether centriole reduplication occurs: normal (pattern 1), precocious centriole disengagement in mitosis (pattern 2), precocious centriole disengagement in the interphase without (pattern 3) and with centriole reduplication (pattern 4). To track centrioles and chromosomes in the mitotic cells, we performed live-imaging using the HeLa-GFP-centrin1 cell line and SiR-DNA, and quantified the frequency of the phenotypes in each siRNA condition. In control cells or Cep57L1-depleted cells, two pairs of centrioles were engaged throughout the interphase and until the end of mitosis (pattern 1) (Figures 6A, 6B and S7A). Single depletion of Cep57 caused precocious centriole disengagement only in mitosis, but not in interphase (pattern 2) (Figures 6A and 6B). As we mentioned above, most Cep57/Cep57L1-depleted cells exhibited precocious centriole disengagement in the interphase (pattern 3), which was frequently followed by centriole reduplication (pattern 4) (Figures 6A and 6B). These phenotypic patterns were consistent with what we had observed in the fixed cells (Figures S7B and S7C). We next quantified the frequency of chromosome segregation errors in each siRNA condition. Cep57- or Cep57/Cep57L1-depleted cells exhibited chromosome segregation errors, which were associated with chromosome misalignment and multipolar spindle formation (siControl; 0.7% ± 1.2%, siCep57; 15.3 ± 4.2%, siCep57/Cep57L1; 30.0 ± 2.0%, N=30, from three independent experiments) (Figures 6C, 6D and Supplementary Movies 3, 4, 5, 6). The frequency of chromosome segregation errors in Cep57/Cep57L1-depleted cells was about twice that in Cep57-depleted cells. To further investigate what factors most effectively led to the increase of the chromosome segregation errors in Cep57/Cep57L1-depleted cells, we next compared the frequency of chromosome segregation errors according to the phenotypic patterns. We revealed that Cep57/Cep57L1-depleted cells with precocious disengagement and centriole reduplication (pattern 4) showed the highest frequency of chromosome segregation errors (57.5 ± 8.0%) (Figures 6C, 6E, S7D and S7E). On the other hand, the frequency of chromosome segregation errors between the cells with precocious disengagement in mitosis (pattern 2, in Cep57-depleted cells, 29.0 ± 8.0%) and interphase (pattern 3, in Cep57/Cep57L1-depleted cells, 34.8 ± 11.6%) were not significantly different (Figure 6E). These results suggest that, upon Cep57 and Cep57L1 depletion, the centriole reduplication induced by precocious centriole disengagement in the interphase is a more direct cause of chromosome segregation errors than the precocious centriole disengagement itself.

**Figure 6.**
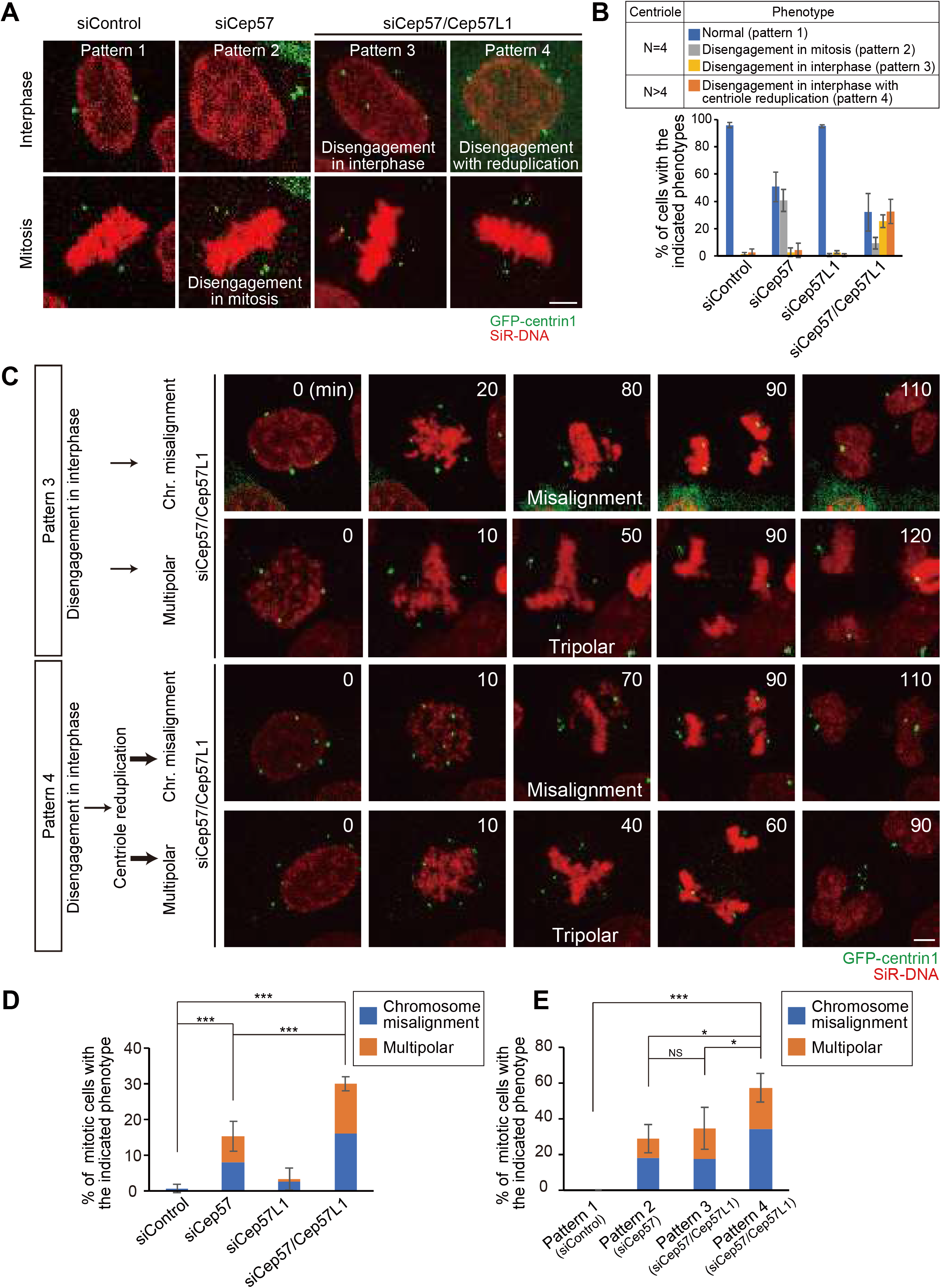
Amplified centrioles caused by co-depletion of Cep57 and Cep57L1 result in high frequency of chromosome segregation errors. (**A**) Phenotypic patterns were categorized into four groups: normal (pattern 1), precocious centriole disengagement in mitosis (pattern 2), precocious centriole disengagement in the interphase without (pattern 3) and with centriole reduplication (pattern 4). HeLa-GFP-centrin1 cells were treated with siControl, siCep57 or siCep57/Cep57L1 in the presence of SiR-DNA (200 nM). (**B**) Histograms represent frequency of mitotic cells with the indicated phenotypes in (A). Values are mean percentages ± s.d. from three independent experiments (n = 50 for each experiment). (**C**) Chromosome segregation errors observed in Cep57/Cep57L1-depleted cells with the indicated phenotype. HeLa-GFP-centrin1 cells were treated with siCep57/Cep57L1 in the presence of SiR-DNA (200 nM). (**D, E**) Histograms represent frequency of the mitotic cells with the indicated chromosome segregation errors observed in (C). Values are mean percentages ± s.d. from three independent experiments (n = 50 for each experiment in (D))(Normal (Pattern 1, siControl): n = 48, 47, 49, Disengagement in mitosis (Pattern 2, siCep57): n = 25, 18, 18, Disengagement in interphase (Pattern 3, siCep57/Cep57L1): n= 14, 14, 10, Disengagement in interphase with reduplication (Pattern 4, siCep57/Cep57L1): n = 21, 16, 12 in (E)). All scale bars, 5 μm. Tukey’s multiple comparisons test was used in (D) and (E) against total value of chromosome misalignment and multipolar to obtain p value. ***p < 0.001; *p < 0.05; NS not significantly different (p > 0.05).

## DISCUSSION

In conclusion, our work is the first to identify Cep57 and Cep57L1 as essential factors maintaining centriole engagement in the interphase. In this study, we also show that this tight regulation of centriole engagement is critical for a proper centriole duplication cycle and chromosome segregation (Figure 7). Surprisingly, depletion of Cep57 and Cep57L1 induced precocious centriole disengagement in the interphase independent of Plk1 activity or cell-cycle arrest. Consistent with previous studies (Lončarek et al., 2010), precocious centriole disengagement in the interphase released mother centrioles from a block to reduplication, and thereby resulted in an increase in the number of centrioles. Furthermore, the number of centrioles per cell increased with each cell division because of the continuous centriole reduplication and inevitable inheritance of the amplified centrioles by daughter cells. In addition, Cep57/Cep57L1-depleted cells exhibited a higher frequency of multipolar spindle formation and chromosome instability mainly because of the amplified centrioles.

**Figure 7.**
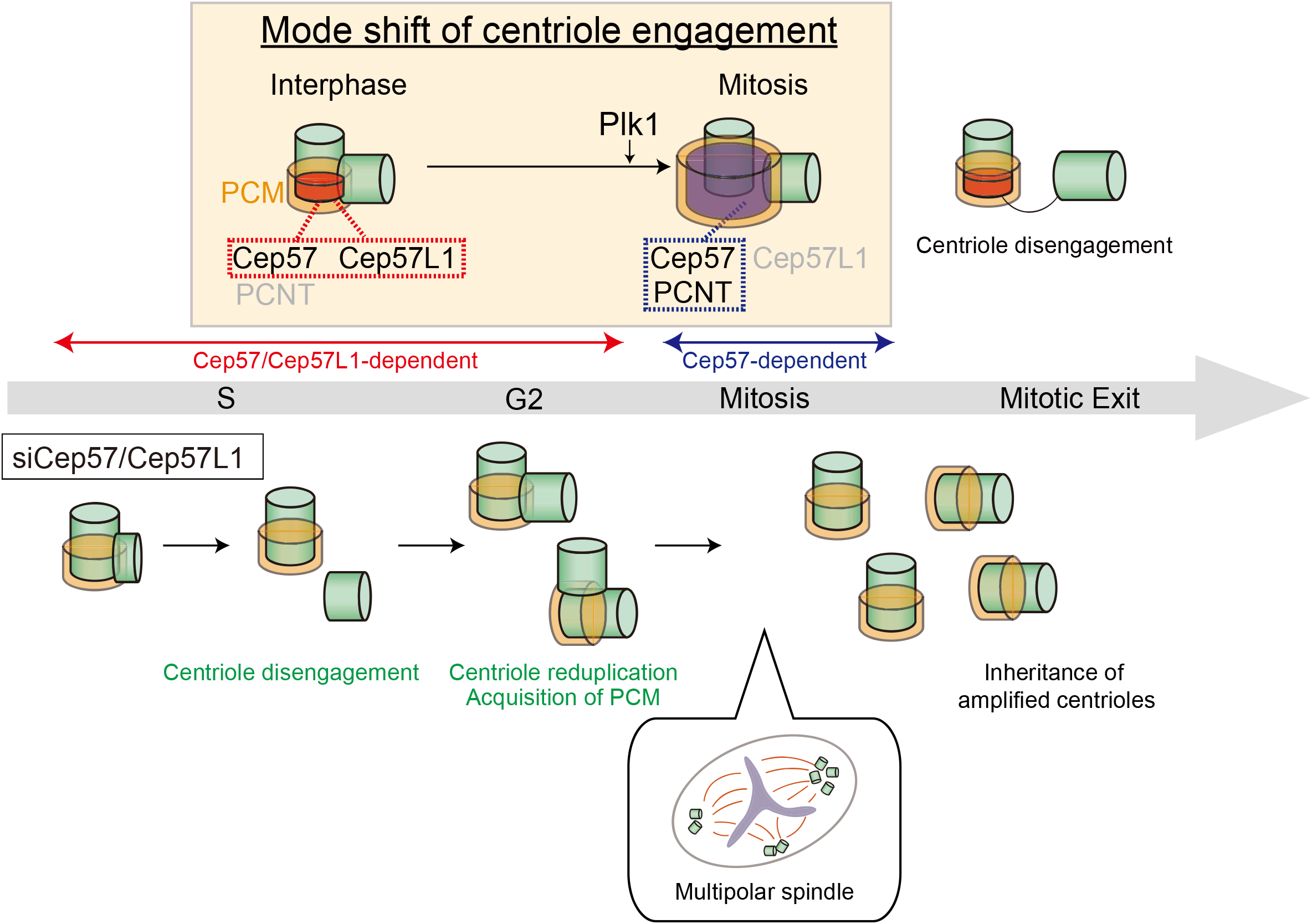
A speculative model for the mode shift of centriole engagement. Centriole engagement in interphase is maintained cooperatively by Cep57 and Cep57L1, and Plk1 changes the engagement mode to Cep57- and PCNT-dependent one at the mitotic entry. Cep57/Cep57L1 co-depletion induces precocious centriole disengagement during interphase, and such disengaged daughter centrioles are converted into centrosomes and can reduplicate before entering mitosis. Consequently, the centriole number increases, which results in the high frequency of chromosome segregation errors.

Although Plk1 is critical for centriole disengagement during mitosis in normal cells (Kim et al., 2015; Tsou et al., 2009), precocious centriole disengagement induced by co-depletion of Cep57/Cep57L1 did not depend on Plk1 activity (Figures 3C and 3D). In addition to Plk1, APC/C and separase are also required for canonical centriole disengagement (Tsou et al., 2009). Separase is a protease activated by APC/C at anaphase onset, and separase cleaves PCNT, a major PCM component, to induce PCM disassembly and centriole disengagement(Lee and Rhee, 2012; Matsuo et al., 2012). Since the artificial activation of APC/C or separase causes centriole disengagement, even in the interphase (Hatano and Sluder, 2012; Prosser et al., 2012), it is possible that co-depletion of Cep57 and Cep57L1 aberrantly activated APC/C or separase in the interphase and thereby caused precocious centriole disengagement. However, even precocious centriole disengagement induced by overexpression of APC/C or separase is suppressed by BI2536 (Prosser et al., 2012). Considering our result showing that Plk1 inhibition did not suppress the phenotype induced by Cep57/Cep57L1 depletion (Figures 3C and 3D), we speculate that co-depletion of Cep57 and Cep57L1 induces precocious centriole disengagement without activating the canonical pathway involving Plk1, APC/C, and separase.

One interesting issue raised in this study is the difference in the timing of precocious centriole disengagement: co-depletion of Cep57/Cep57L1 causes the phenotype in the interphase whereas a single depletion of Cep57 causes the phenotype only in mitosis (Figure 1) (Watanabe et al., 2019). Similar to Cep57, PCNT is involved in the maintenance of centriole engagement only in mitosis, but the depletion of PCNT does not disrupt centriole engagement in the interphase (Figure 1). These results imply that the mode of centriole engagement somehow changes from interphase to mitosis. Importantly, the surrounding PCM expands dynamically and functionally matures toward mitosis (Khodjakov and Rieder, 1999), which is likely a critical event for switching the status of centriole engagement. This PCM maturation at the G2/M transition is known to be a Plk1-dependent process, which includes dynamic reorganization of the PCM from interphase (highly ordered state) to mitosis (amorphous state) (Lawo et al., 2012). In addition, a recent study showed that the activity of Plk1 is needed for extending the distance between mother and daughter centrioles from 50 nm to 80 nm in early mitosis (Shukla et al., 2015). Given these observations, we speculate that the Plk1-dependent dynamic rearrangement of PCM components couples the PCM expansion with changes in the status of centriole engagement. In line with this idea, the precocious centriole disengagement was induced only in mitosis by Cep57 depletion, and this could be suppressed by treatment with a Plk1 inhibitor (BI2536) (Watanabe et al., 2019; Wilhelm et al., 2019). Meanwhile, the precocious centriole disengagement that occurred in the interphase after co-depletion of Cep57 and Cep57L1 was not suppressed by Plk1 inhibition. It has also been reported that Plk1-dependent phosphorylation of PCNT is required for PCM expansion (Lee and Rhee, 2011). However, how Plk1 and PCNT cooperatively regulate PCM expansion and the shift of the centriole engagement mode in early mitosis remains elusive.

According to the findings in this study, the timing of centriole disengagement in the cell cycle is also critical for the occurrence of centriole reduplication. Given that precocious centriole disengagement in Cep57-depleted cells was not accompanied by centriole reduplication, the reloading of centriole duplication factors must be tightly restricted in mitosis so as not to increase the centrosome number. In contrast, in Cep57/Cep57L1-depleted cells, precocious centriole disengagement occurs in interphase, which secures sufficient centriole components and enough time for centriole duplication to occur. Such centriole reduplication thereby results in the drastic increase of centrosome numbers, which then more frequently leads to chromosome segregation errors. Thus, on the basis of these observations, it is conceivable that the tight control of maintenance of centriole engagement is more important in interphase than in mitosis

**Supplementary Figure 1.** (**A**) Alignments of full-length *H. sapiens* Cep57L1 and Cep57. Asterisks indicate the residues identical in aligned sequence; colons: conserved substitutions; periods: semi-conserved substitutions. The position of PINC motif and predicted microtubule binding domain are indicated in pink and green boxes. (**B**) The phenotype induced by Cep57/Cep57L1 co-depletion was confirmed by using another siRNA. HeLa cells were treated with siCep57 and siCep57L1#2, and immunostained with antibodies against centrin2 (green) and Cep192 (red). siCep57L1#2 targets a different sequence in open reading frame from siCep57L1#1 which is used in main figures. (**C**) U2OS cells also exhibited precocious centriole disengagement and amplified centrioles upon Cep57/Cep57L1 co-depletion. U2OS cells were treated with siControl or siCep57/Cep57L1, and immunostained with antibodies against centrin2 (green) and Cep192 (red). All scale bars, 5 μm in the low-magnified view, 1 μm in the inset.

**Supplementary Figure 2.** (**A**) The number of old mother centriole was one irrespective of the phenotypes in Cep57/Cep57L1-depleted cells. HeLa cells were treated with siControl or siCep57/Cep57L1, and immunostained with antibodies against centrin2 (green) and ODF2 (red). Scale bars, 5 μm in the low-magnified view, 1 μm in the inset (**B**) Histograms represent frequency of the interphase cells with the indicated number of ODF2 foci observed in (A). Values are mean percentages ± s.d. from two independent experiments (n = 50 for each experiment). (**C**) Quantification of the duration from anaphase onset to next mitotic entry. HeLa-GFP-centrin1 cells were treated with siControl or siCep57/Cep57L1, and visualized for live imaging for 48 h (n = 25). (**D**) Cep57/Cep57L1 co-depletion did not affect cell cycle progression. HeLa cells were treated with the indicated siRNAs, and followed by flow cytometry analysis. Two-tailed, unpaired Welch’s t-test was used in (C) to obtain p value. NS not significantly different (p > 0.05).

**Supplementary Figure 3.** (**A**) POC5 was recruited to a portion of disengaged centrioles. HeLa cells were treated with siControl or siCep57/Cep57L1, and immunostained with antibodies against centrin2 (green) and POC5 (red). (**B**) Histograms represent frequency of cells with the indicated number of POC5 positive centrioles among cells with two pairs of centrioles or disengaged four centrioles observed in (A). Values are mean percentages ± s.d. from three independent experiments (n = 30 for each experiment). (**C**) Cep295 was recruited to the disengaged centrioles. HeLa cells were treated with siControl or siCep57/Cep57L1, and immunostained with antibodies against centrin2 (green) and Cep295 (red). Scale bars, 5 μm in the low-magnified view, 1 μm in the inset.

**Supplementary Figure 4.** (**A**) PCNT was recruited to disengaged daughter centrioles mainly in the G2 phase and mitosis. HeLa-GFP-centrin1 cells were treated with siCep57/Cep57L1, and immunostained with antibodies against GFP (green), PCNT (red) and CENP-F (cyan). White/black arrowheads indicate PCNT positive/negative centrioles, respectively. Scale bar, 5 μm. (**B**) Histograms represent frequency of cells with >2 PCNT positive centrioles among cells with disengaged four centrioles before, in and after the G2 phase observed in (A). Values are mean percentages ± s.d. from three independent experiments (n = 30 for each experiment). (**C**) Schematic illustration of the result in (A) and (B). PCNT was recruited onto disengaged daughter centrioles mainly in the G2 phase and mitosis. (**D**) Ectopic MTOC activity of precociously-disengaged daughter centrioles in Cep57/Cep57L1-depleted cells. HeLa cells were treated with siControl or siCep57/Cep57L1 and followed by nocodazole treatment (10 μM) for 3 h. After nocodazole treatment, the cells were cold-treated for 1 h, followed by 1 min incubation at 37°C and immunostaining with antibodies against EB1 (green) and CP110 (red). White arrowheads indicate centrosomes. Scale bar, 5 μm.

**Supplementary Figure 5.** (**A**) The Cep57L1 PINC motif was not required for the centrosome localization. HeLa cells expressing FLAG-Cep57L1 or FLAG-Cep57L1 mutant lacking the PINC motif (Δ52-86 a.a.) were immunostained with antibodies against FLAG (green) and Cep192 (red). (**B**) The Cep57L1 PINC motif was not required for binding to the C-terminal region of PCNT containing the conserved PACT domain. HEK293T cells co-expressing FLAG empty (control), FLAG-Cep57, FLAG-Cep57L1 or FLAG-Cep57L1 mutant lacking the PINC motif and GFP-PCNT (3132-3336 a.a.) were immunoprecipitated with FLAG antibodies and immunoblotted with the indicated antibodies. (**C**) Overexpressed Cep57 was accumulated on the microtubules, but overexpressed Cep57L1 aggregated in the cytoplasm. HeLa cells expressing HA-Cep57 or HA-Cep57L1 were immunostained with antibodies against HA (red) and α-tubulin (green). (**D**) Cep57L1 formed homodimer. HEK293T cells co-expressing FLAG empty (control), FLAG-Cep57 or FLAG-Cep57L1 and HA-Cep57 or HA-Cep57L1 were immunoprecipitated with FLAG antibodies and immunoblotted with the indicated antibodies. All scale bars, 5 μm in the low-magnified view, 1 μm in the inset.

**Supplementary Figure 6.** (**A**) The centrosome localization of Cep57 and Cep57L1 was dependent on Cep63 and Cep152. HeLa cells were treated with siControl, siCep63 or siCep152, and immunostained with antibodies against Cep57 (green), Cep57L1 (red) and Cep192 (cyan). (**B**) The centrosome localization of Cep63 was partially dependent on Cep57 and Cep57L1. HeLa cells were treated with siControl or siCep57/Cep57L1, and immunostained with antibodies against ODF2 (green), Cep63 (red) and GT335 (cyan). (**C**) The centrosome localization of Cep152 was partially dependent on Cep57 and Cep57L1. HeLa cells were treated with siControl or siCep57/Cep5L1, and immunostained with antibodies against ODF2 (green), Cep152 (red) and GT335 (cyan). All scale bars, 5 μm in the low-magnified view, 1 μm in the inset.

**Supplementary Figure 7.** (**A**) Normal bipolar spindle formation (pattern 1) was observed in Cep57L1-depleted cells. HeLa-GFP-centrin1 cells were treated with siCep57L1 in the presence of SiR-DNA (200 nM). Scale bar, 5 μm. (**B**) Phenotypic patterns observed in fixed cells are consistent with live-imaging analysis. HeLa cells were treated with siControl, siCep57, siCep57L1 or siCep57/Cep57L1, and immunostained with antibodies against centrin2 (green) and Cep192 (red). Scale bars, 5 μm in the low-magnified view, 1 μm in the inset. (**C**) Histograms represent frequency of mitotic cells with the indicated phenotypes observed in (B). Values are mean percentages ± s.d. from three independent experiments (n = 30 for each experiment). (**D**) Normal bipolar spindle formation observed in control and Cep57L1-depleted cells. HeLa-GFP-centrin1 cells were treated with siControl or siCep57L1 in the presence of SiR-DNA (200 nM). Scale bar, 5 μm. (**E**) Chromosome segregation errors observed in Cep57-depleted cells. HeLa-GFP-centrin1 cells were treated with siCep57 in the presence of SiR-DNA (200 nM). Scale bar, 5 μm.

### MATERIALS AND METHODS

#### Cell culture and transfection

HeLa, U2OS and HEK293T cells were obtained from the ECACC (European collection of cell cultures). HeLa, U2OS and HEK293T cells were cultured in Dulbecco’s modified Eagle’s medium (DMEM) supplemented 10% fetal bovine serum (FBS) and 100 μg/ml penicillin-streptomycin at 37◻°C in a 5% CO_2_ atmosphere. Transfection of siRNA or DNA constructs into HeLa and HEK293T cells was conducted using Lipofectamine RNAiMAX (Life Technologies) or Lipofectamine 2000 (Life Technologies), respectively. Unless otherwise noted, the transfected cells were analyzed 48◻h after transfection with siRNA and 24h after transfection with DNA constructs. When the cells were analyzed 96 h after transfection with siRNA, additional siRNA was transfected 48 h after the first transfection

#### RNA interference

The following siRNAs were used: Silencer Select siRNA (Life Technologies) against Cep57 (s18692), Cep57L1#1 (s226224), Cep57L1#2 (s226223), Cep63 (s37123), Cep152 (s225921) and negative control (4390843). Unless otherwise noted, Cep57L1 #1 were used in this study

#### Plasmids

Complementary DNA (cDNA) encoding Cep57L1 isoform 1 (NCBI NP_001258781.1) was amplified from cDNA library of A549 cells. The Cep57L1 cDNA was subcloned into pCMV5-HA (Addgene) and pCMV5-FLAG (Addgene). pCMV5 constructs encoding full-length Cep57 and pTB701 constructs encoding PCNT were described previously (Watanabe et al., 2019; Takahashi et al., 2002). The Cep57L1 deletion mutant constructs were created using PrimeSTAR mutagenesis basal kit (TaKaRa) and In-Fusion HD cloning kit (Clontech) according to manufacturer’s protocol

#### Antibodies

The following primary antibodies were used in this study: rabbit antibodies against Cep57L1 (Proteintech, 24957-1-AP, IF 1:500), Cep63 (Proteintech, 16268-1-AP, IF 1:1000), PCNT (Abcam, ab4448, IF 1:2000), Cep192 (Bethyl Laboratories, A302–324A, IF 1:1000), Cep152 (Bethyl Laboratories, A302–480A, IF 1:1000), CP110 (Proteintech, 12780–1-AP, IF 1:500), ODF-2 (Abcam, ab43840, IF 1:1000), GFP (MBL, 598, WB 1:1000), Cep295 (Merck, HPA038596, IF 1:500), CENP-F (Abcam, ab108483, IF 1:500), POC5 (Bethyl Laboratories, A303-341A, IF 1:1000), FLAG-tag (Merck, F7425, IF 1:1000, WB 1:1000) and HA-tag (Abcam, ab9110, IF 1:1000, WB 1:1000); mouse antibodies against Cep57 (Abcam, ab169301, IF1:1000), PCNA (Santa Cruz Biotechnology, sc-56, IF 1:1,000), PCNT (Abcam, ab28144, IF 1:1000), centrin2 (Merck, 20H5, IF 1:500), EB1 (BD Transduction Laboratories, 610534, IF 1:1000), HsSAS-6 (Santa Cruz Biotechnology, sc-81431, IF 1:300), Polyglutamylation Modification (GT335) (AdipoGen, AG-20-B0020-C100, IF 1:2000), γ-tubulin (GTU88) (Sigma-Aldrich, T5192, IF 1:1000), FLAG-tag (Merck, F3165, IF 1:1000, WB 1:1000) and α-tubulin (Merck, DM1A, IF1:1000, WB 1:1000); goat antibody against GFP (Abcam, ab6662, IF 1:500, conjugated to FITC); Alexa 488-labelled ODF-2 (Abcam, ab43840, IF 1:500) and Alexa 647-labelled Cep192 (Bethyl Laboratories, A302–324A, IF 1:500) were generated with Alexa Fluor labelling kits (Life Technologies). The following secondary antibodies were used: Alexa Fluor 488 goat anti-mouse IgG (H+L) (Molecular Probes, A-11001, 1:1000), Alexa Fluor 647 goat anti-mouse IgG (H+L) (Abcam, ab150115, 1:1000), Alexa Fluor 568 goat anti-rabbit IgG (H+L) (Molecular Probes, A-11011, 1:1000) for IF; Goat polyclonal antibodies-horseradish peroxidase against mouse IgG (Promega, W402B, 1:10000) and rabbit IgG (Promega, W401B, 1:10000) for WB

#### Chemicals

The following chemicals were used in this study: nocodazole (Wako, 31430-18-9), RO3306 (SIGMA, SML0569), BI2536 (AdooQ, A10134) and SiR-DNA (Spirochrome, CY-SC007)

#### Microscopy

For immunofluorescence analysis, the cells cultured on coverslips (Matsunami, 18 mm, thickness No. 1_0.12-0.17 mm for widefield microscope) were fixed using −20◻°C methanol for 7◻min and washed with PBS. The cells were permeabilized after fixation with PBS/0.05% Triton X-100 (PBSX) for 5◻min twice, and incubated for blocking in 1% BSA in PBSX for 30◻min at room temperature (RT). The cells were incubated with primary antibodies for 24◻h at 4◻°C or 2 h at RT, washed with PBSX three times, and incubated with secondary antibodies for 1◻h at RT. The cells were thereafter washed with PBSX twice, stained with 0.2◻μg/ml Hoechst 33258 (DOJINDO) in PBS for 5◻min at RT, washed again with PBSX and mounted onto glass slides.

Taking the images and counting the number of immunofluorescence signals were preformed using an Axioplan2 fluorescence microscope (Carl Zeiss) with 63 × or 100 × /1.4 NA plan-APOCHROMAT objectives and DeltaVision Personal DV-SoftWoRx system (Applied Precision) equipped with a CoolSNAP CH350 CCD camera.

SIM images were taken by Elyra 7 (Carl Zeiss Microscopy GmbH, Jena, Germany) equipped with a x63 ZEISS alpha Plan-Apochromat N.A. 1.46 objective and PCO edge sCMOS camera

#### Live imaging

Cell Voyager CQ1 (Yokogawa Electric Corp) equipped with a 40 × dry objective lens was used for live cell imaging. HeLa cells stably expressing GFP-centrin1 were cultured on 24-well glass bottom plate (Greiner-bio-one, #662892) at 37◻°C in a 5% CO_2_ atmosphere. Before imaging, cells were treated with siRNAs for 24 h or 72h and with 200 nM of SiR-DNA for 6 h. Images were taken by sCMOS camera. After 24 or 72 h from transfection, the cells were visualized every 10◻min over 24 h or 48 h. The images were collected at 1.2 μm z steps. Maximum projections were generated using ImageJ (National Institutes of Health).

#### Immunoprecipitation and western blotting

For preparation of human cell lysates for western blotting, HEK293T cells were collected 24 h after transfection, lysed on ice in lysis buffer (20◻mM Tris/HCl pH7.5, 50 mM NaCl, 1% TritonX-100, 5◻mM EGTA, 1◻mM DTT, 2◻mM MgCl_2_ and 1/1000 protease inhibitor cocktail (Nakarai tesque)). Insoluble material was removed after centrifugation for 10◻min. For IP of FLAG-tagged proteins, whole cell lysates were incubated with FLAG antibody-conjugated M2 agarose gel (Merck) for 2◻h at 4◻°C. The beads were washed at least three times with lysis buffer and resuspended in SDS-sample buffer before loading onto 8% polyacrylamide gels, followed by transfer on Immobilon-P membrane (Merk). The membrane was probed with the primary antibodies, followed by incubation with their respective horseradish peroxidase-conjugated secondary antibodies (Promega). The membrane was soaked with Amersham ECL Prime (GE Healthcare) or Chemi-Lumi One Ultra (Nakarai tesque). Washes were performed in PBS containing 0.02% Tween-20 (PBST). The signal was detected with Chemi Doc XRS+ (BIO RAD). Unless otherwise specified, the experiments of western blotting were repeated at least three times. The antibody against α-tubulin was used as a loading control

## Data availability

The data that support the findings of this study are available from the corresponding author upon request

## Contribution

K.K.I., K.W. and D.K. designed the study; K.K.I., K.W., T.C., S.H. and D.K. designed experiments; K.K.I., K.W., H.I. and K.M. performed experiments; K.K.I. and H.I analyzed data; K.K.I., K.W. and D.K. wrote the manuscript, which was commented on by all authors.

## Acknowledgements

We gratefully acknowledge A. Sekigawa for technical support in SIM microscopy; and the members of the Kitagawa laboratory for technical support and discussion. This work was supported by a Grant-in-Aid for Scientific Research (S) and Research Activity start-up from the Ministry of Education, Science, Sports and Culture of Japan, by the Takeda Science Foundation, by the Japan Science Society (The Sasagawa Scientific Research Grant), by the Daiichi Sankyo Foundation of Life Science.

