## Supplementary figures for "Cep57 and Cep57L1 cooperatively maintain centriole engagement during interphase to ensure proper centriole duplication cycle"

### Supplementary Figure 1

**A**

PINC motif (highlighted: Cep57L1 residues 59–89, Cep57 residues 73–103)

Predicted MT binding domain (highlighted: Cep57L1 residues 313–363, Cep57 residues 350–400)

```
Cep57L1  1    MD------------------------SELMHSIVGSYHKPPERVFVPSFTQNEPSQNCHPANLEVTSPKILHSPNSQALILALKTLQEKIHRLE  70
  Cep57  1    MAAASVSAASGSHLSNSFAEPSRSNGSMVRHSSSPYVVYPSDKPFLNSDLRRSPS----------KPTLAYPESNSRAIFSALKNLQDKIRRLE  84
              *                         * : **       * ::*: *  :..**          .      . **:*:: ***.**:**:***

Cep57L1  71   LERTQAEDNLNILSREAAQYKKALENETNERNLAHQELIKQKKDISIQLSSAQSRCTLLEKQLEYTKRMVLNVEREKNMILEQQAQLQREKEQD  164
  Cep57  85   LERIQAEESVKTLSRETIEYKKVLDEQIQERENSKNEESKHNQELTSQLLAAENKCNLLEKQLEYMRNMIKHAEMERTSVLEKQVSLERERQHD  178
              *** ***:.:: ****: :***.*::: :**: :::*  *:::::: ** :*:.:*.******** :.*: :.* *:. :**:*..*:**:::*

Cep57L1  165  QMKLYAKLEKLDVLEKECFRLTTTQKTAEDKIKHLEEKLKEEEHQRKLFQDKASELQTGLEISKIIMSSVSNLKHSK-----EKKK----SSKK  249
  Cep57  179  QTHVQSQLEKLDLLEQEYNKLTTMQALAEKKMQELEAKLHEEEQERKRMQAKAAELQTGLETNRLIFEDKATPCVPNARRIKKKKSKPPEKKSS  272
              * :: ::*****:**:*  :*** *  **.*::.** **:***::** :* **:******* .::*:.. :.    :     :**.    ....

Cep57L1  250  TKCIKRRPPWQICSKFGALPFVAEKMRQHRDPHILQKPFNV------TETRCL-------PKPSR---TTSWCKAIPPDSEKSISICDNLSELL  327
  Cep57  273  RNYFGAQPHYRLC--LGDMPFVAGKSTSPSHAVVANVQLVLHLMKQHSKALCNDRVINSIPLAKQVSSRGGKSKKLSVTPPSSNGINEELSEVL  364
               : :  :* :::*  :* :**** *   .  . : :  : :      ::: *        *  .:     . .* :     .* .* ::***:*

Cep57L1  328  MAMQDELDQMSMEHQELLKQMKETESHSVCDDIECELECLLKKMEIKGEQISKLKKHQDSVCKLQQKVQNSKMSEASGIQQEDSYPKGSKNIKN  421
  Cep57  365  QTLQDEFGQMSFDHQQLAKLIQESPTVELKDKLECELEALVGRMEAKANQITKVRKYQAQLEKQKLEKQKKELKATKKTLDEERNSSSRSGITG  458
               ::***:.***::**:* * ::*: : .: *.:*****.*: :** *.:**:*::*:* .: * : : *:.::. :.    :*:   .. ..*..

Cep57L1  422  SP-RKCLT---DTNLFQKNSSFHPIRVHNLQMKLRRDDIMWEQ  460
  Cep57  459  TTNKKDFMKLRPGEKRRKN-LQLLKDMQSIQNSLQSSSLCWDY  500
              :  :* :      :  :**       ::.:* .*: ..: *:
```

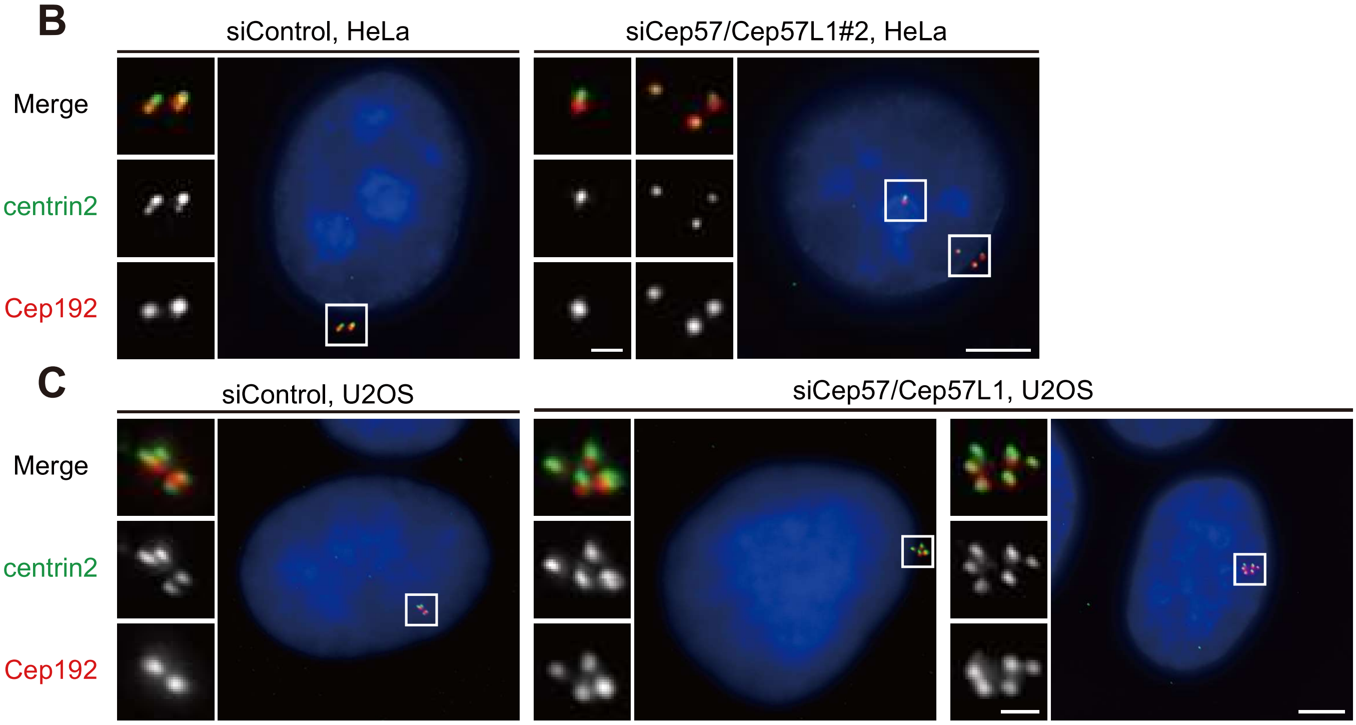

Supplementary Figure 2

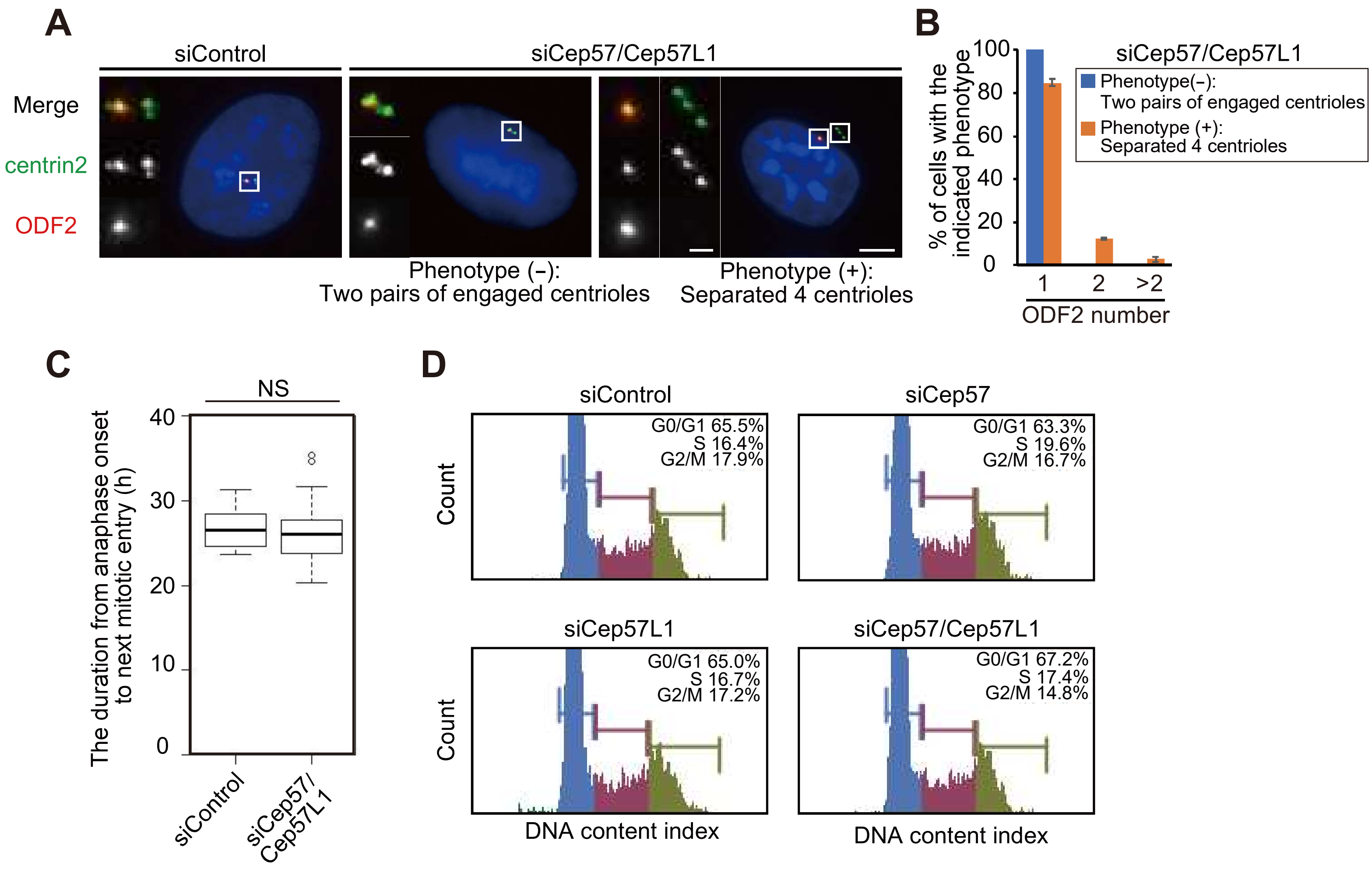

### Supplementary Figure 3

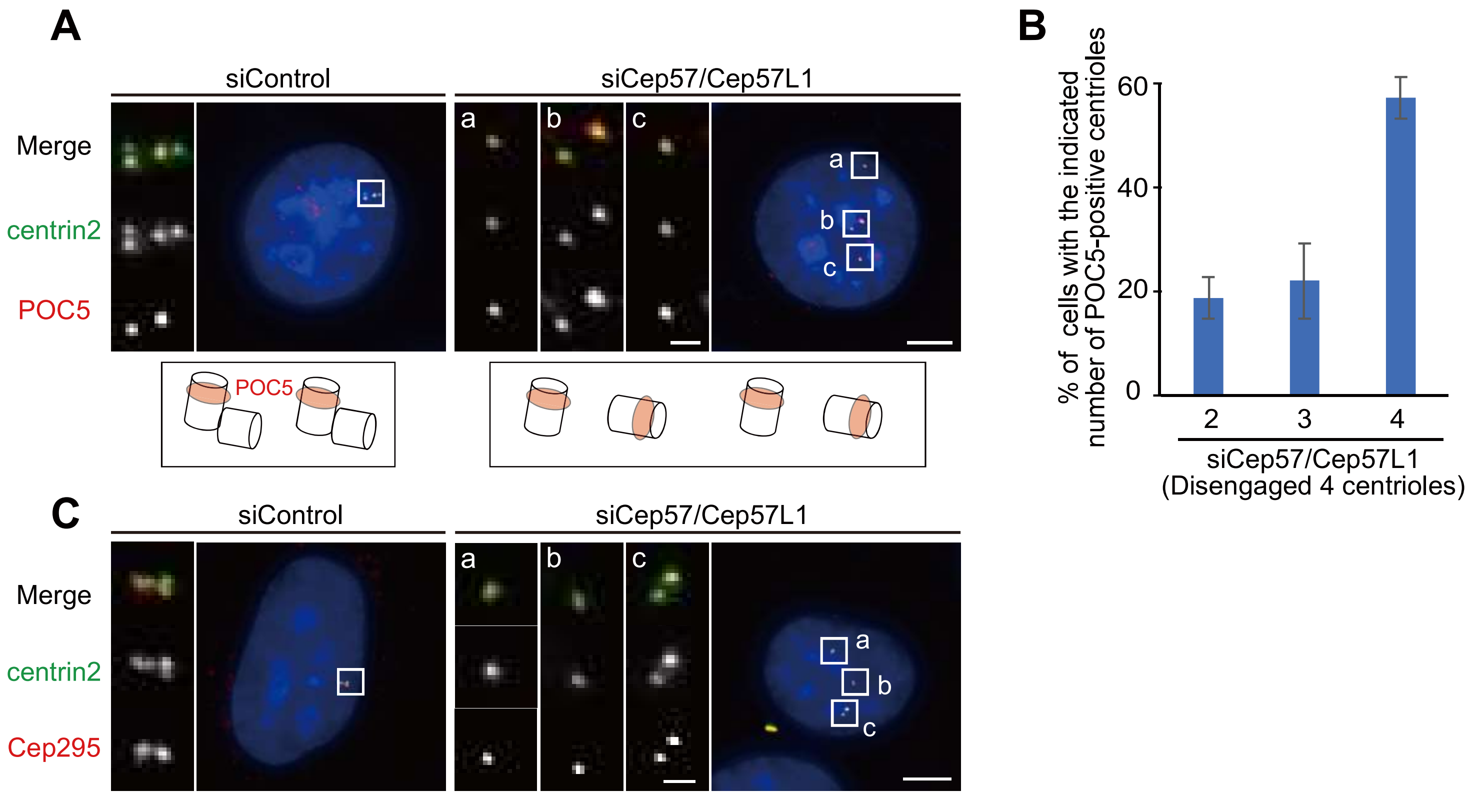

### Supplementary Figure 4

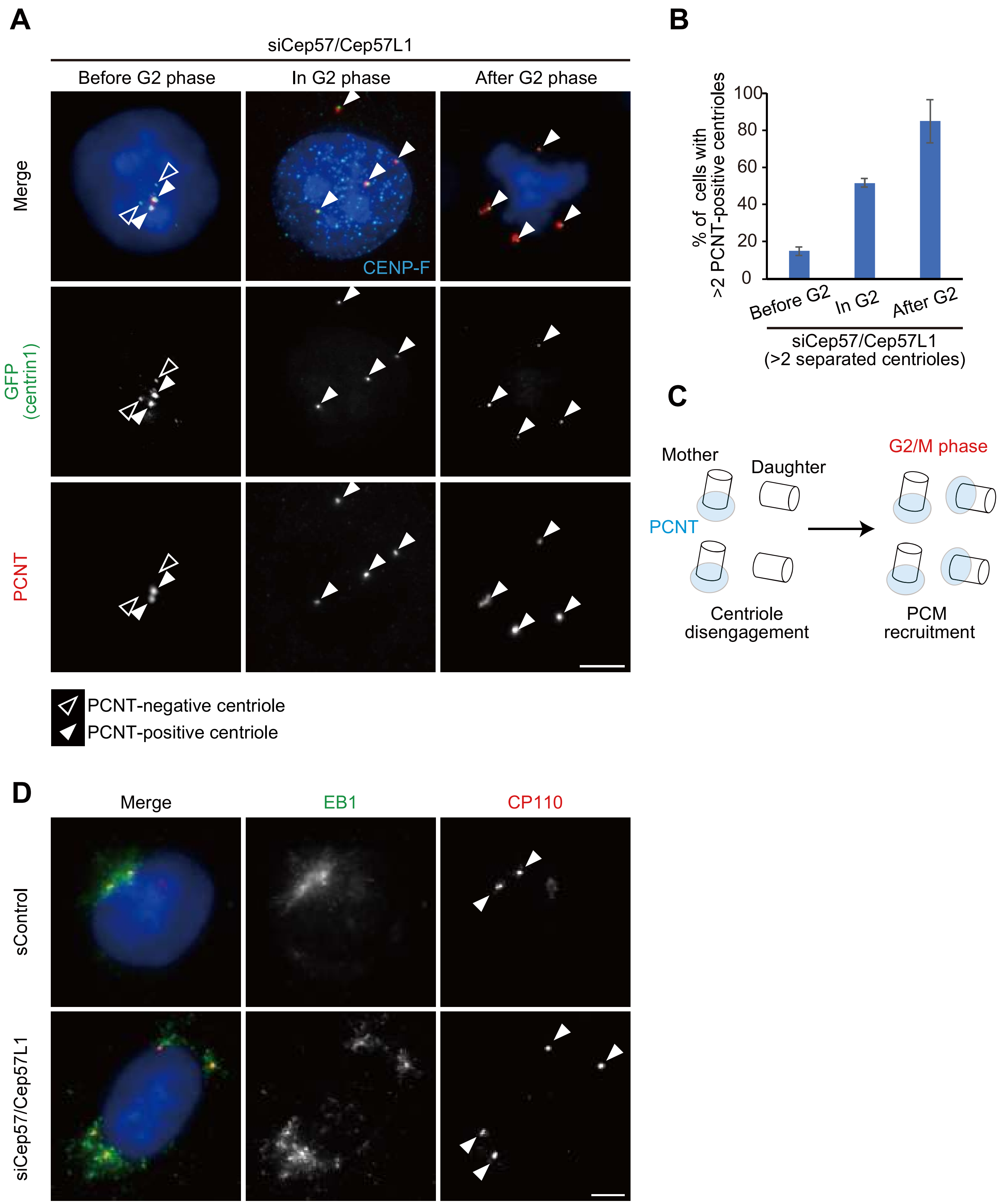

### Supplementary Figure 5

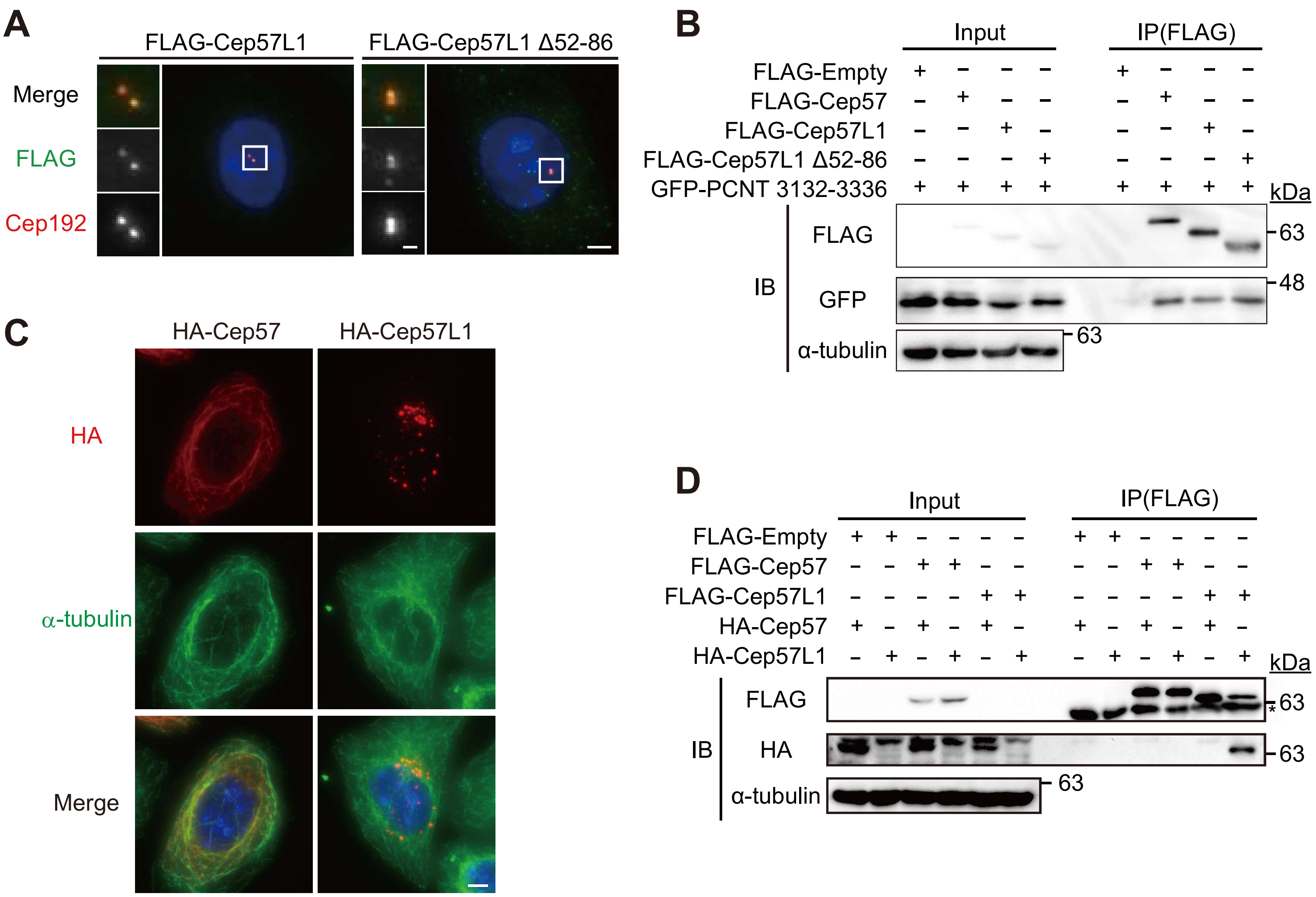

Supplementary Figure 6

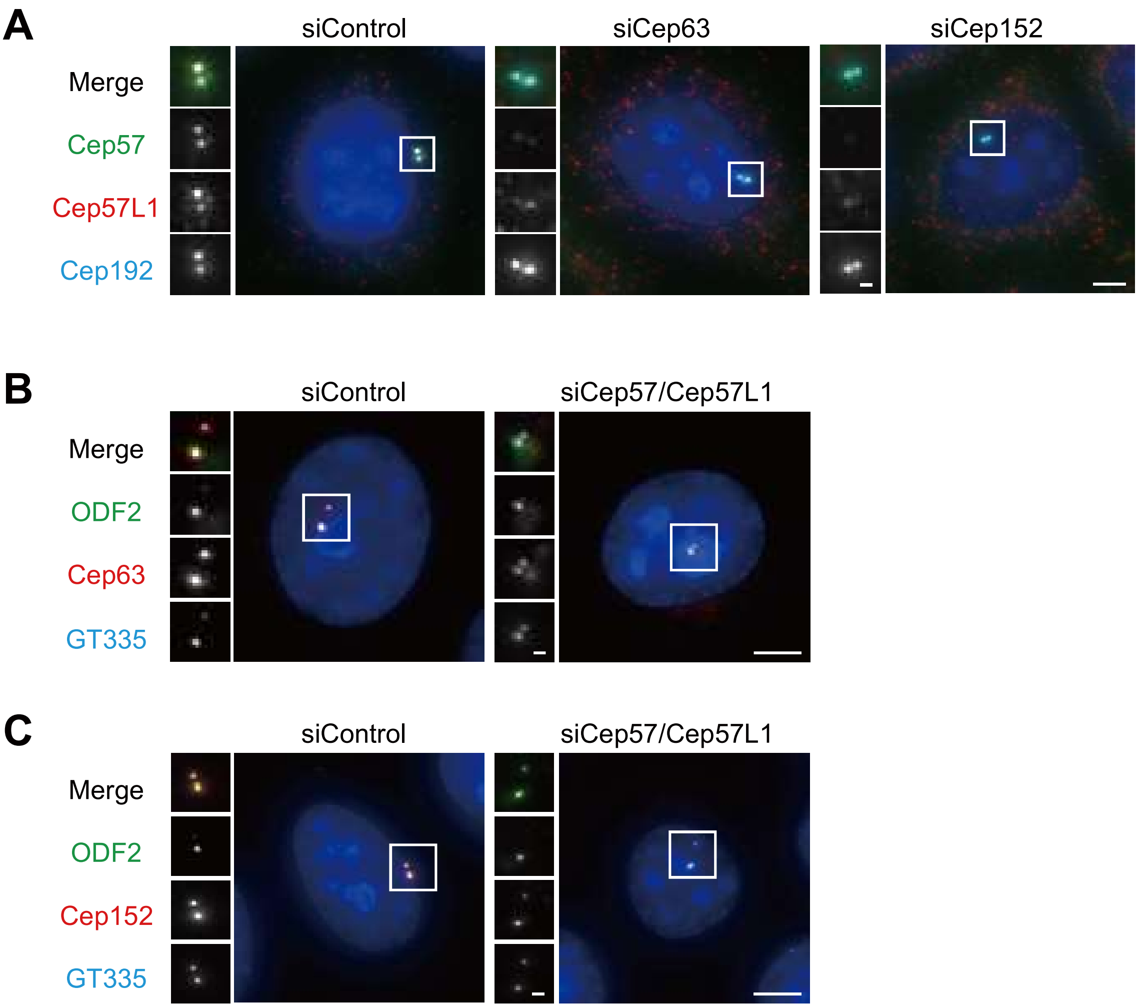

### Supplementary Figure 7

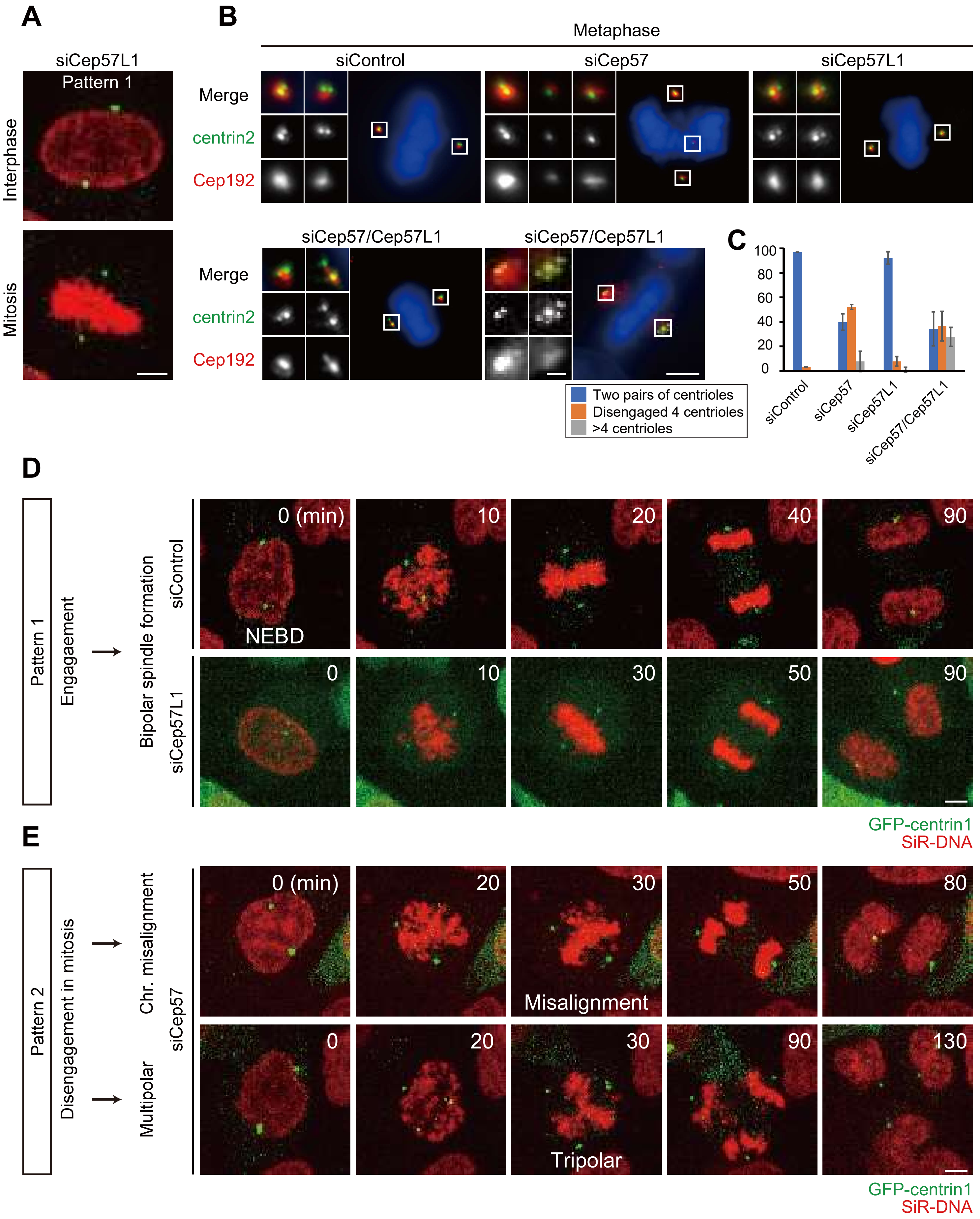
